# Role of spinal sensorimotor circuits in triphasic command: a simulation approach using Goal Exploration Process

**DOI:** 10.1101/2023.12.22.572982

**Authors:** Daniel Cattaert, Matthieu Guemann, Florent Paclet, Bryce Chung, Pierre-Yves Oudeyer, Aymar de Rugy

**Author notes:** **Corresponding author:** Daniel Cattaert, INCIA UMR 5287 – 146 rue Léo Saignat, 33076 BORDEAUX CEDEX, France.

## Abstract

During rapid voluntary limb movement about a single joint, a stereotyped triphasic pattern is typically observed in the electromyograms (EMGs) of antagonistic muscles acting at this joint. To explain the origin of such triphasic commands, two types of theories have been proposed. Peripheral theories consider that triphasic commands result from sensorimotor spinal networks, either through a combination of reflexes or through a spinal central pattern generator. Central theories consider that the triphasic command is elaborated in the brain. Although both theories were partially supported by physiological data, there is still no consensus about how exactly triphasic commands are elaborated. Moreover, capacities of simple spinal sensorimotor circuits to elaborate triphasic commands on their own have not been tested yet. In order to test this, we modelled arm musculoskeletal system, muscle activation dynamics, proprioceptive spindle and Golgi afferent activities and spinal sensorimotor circuits. Step commands were designed to modify the activity of spinal neurons and the strength of their synapses, either to prepare (SET) the network before movement onset, or to launch the movement (GO). Since these step commands do not contain any dynamics, changes in muscle activities responsible for arm movement rest entirely upon interactions between the spinal network and the musculo-skeletal system. Critically, we selected step commands using a Goal Exploration Process inspired from baby babbling during development. In this task, the Goal Exploration Process proved very efficient at discovering step commands that enabled spinal circuits to handle a broad spectrum of functional behaviors, displayed in a behavioral space characterized by movement amplitude and maximal speed. All over the behavioral space, specific SET and GO commands elicited natural triphasic commands, thereby substantiating the inherent capacity of the spinal network in generating them.

**Key points Summary:**

- Spinal sensorimotor circuits, despite extensive physiological analyses, remain poorly understood in the context of voluntary movement.
- This study models the capabilities of these spinal networks, alongside musculoskeletal dynamics, to achieve single-joint arm flexion following minimum jerk principles.
- Preparatory (SET) and launch (GO) step commands modulate spinal network states, including neuron activity and synaptic strength.
- Notably, arm movement results from sensorimotor interactions within spinal circuits, even with basic step commands that have no dynamics.
- A Goal Exploration Process (GEP), inspired by developmental baby babbling through trial and error, was used to refine step commands.
- The spinal networks, when in appropriate states, were shown to spontaneously generate triphasic commands across a broad range of valid movement amplitudes and speeds.
- These findings highlight the intrinsic capacity of spinal sensorimotor circuits to manage complex movement dynamics, offering novel testable insights as to their possible functional role in voluntary motor control.

## Introduction

During rapid voluntary limb movement about a single joint, a stereotyped triphasic pattern is observed in the electromyograms (EMGs) of antagonistic muscles commanding this joint (Hallett *et al*., 1975). Prior to limb displacement, the agonist muscle is briefly active (AG1). After the limb begins to move, a burst of muscle activity occurs in the antagonist muscle (ANT) while the agonist is silenced, and then a second burst is observed in the agonist (AG2). The precise organization of this triphasic pattern ensures the smoothness of the movement.

What is the origin of this triphasic command? Two possibilities have been proposed; either a **spinal origin** via sensorimotor circuits (Barnett & Harding, 1955; Terzuolo *et al*., 1974; Ghez & Martin, 1982) or a **central origin** elaborated in brain networks and transmitted to the spinal cord via descending commands. The **central hypothesis** was supported by observations on humans with severe, though not complete, sensory neuropathy, who could still produce triphasic patterns (Hallett *et al*., 1975; Rothwell *et al*., 1982). Although these studies highlight the capacities of the motor cortex and other higher centers that can learn how to control limb movements after partial or total deafferentation, the deep reorganization of motor commands following such deafferentation prevents firm conclusion about how triphasic commands are produced in healthy subjects. Other studies used reversible transient deafferentation using ischemic process of the muscles controlling the joint and their afferent nerves Ia and Ib, the diameter of which are more sensitive to ischemia than axons of alpha motoneurons (Sanes & Jennings, 1984). During such reversible ischemia, the triphasic pattern persists, although passive movements fail to elicit stretch reflexes.

Other experiments indicate that triphasic motor command is also present in the primary motor cortex (M1), and that neural activities in M1 are correlated with output kinematics and kinetics during isometric-force and arm-reaching tasks (Sergio *et al*., 2005). Physiological model describing the transformation from the neural activity in M1, through the muscle control signal, into joint torques and down to endpoint forces and movements (Trainin *et al*., 2007), show that optimized solutions for M1 activities display triphasic changes in temporal pattern and instantaneous directionality similar to that found in the EMG (Trainin *et al*., 2007).

In opposition with the arguments in favor of a central origin of triphasic commands, it was shown that procedures that modify discharge of muscle spindle afferents during movement alter the duration and amplitude of these bursts (Hallett & Marsden, 1979; Brown & Cooke, 1981). In addition, simulation studies concerning the central origin of triphasic commands, although aiming toward physiological models, often lack spinal sensorimotor loops (Trainin *et al*., 2007). Therefore, if we cannot doubt that M1 neurons and populations of neurons present activities correlated with kinetic, kinematic and dynamic cues of the performed movements, there is still questions about the possible role of spinal sensorimotor circuits in the elaboration of triphasic commands. Notably, since both efference copy from spinal interneurons and afferent feedback from proprioceptors project onto sensorimotor cortical areas, they constitute a potential origin for triphasic patterns observed in cortical areas. Consequently, if the first EMG activity (AG1) can be attributed to a central command from the brain (at least for its onset), there is still debate about ANT and AG2.

**The aim of this paper was to explore the capacities of the spinal sensorimotor circuits in shaping triphasic commands**, using a modeling approach based on the neuromechanical environment AnimatLab (https://www.animatlab.com/). We modelled the movement of the forearm around the elbow joint, involving two muscles (biceps and triceps). Movements consisted in elbow flexion starting from a fully extended position. Given our aim, we could not use complex descending commands because if descending commands already contain triphasic patterns it would have been impossible to analyze spinal network circuits capacity to produce a triphasic pattern by their own. Therefore, rather than using descending commands from the brain, we aimed at searching states in which sensorimotor circuits could produce by themselves complex dynamic muscle activities observed during rapid arm flexion in human. Our approach is inspired from spinal like regulator (SLR) work (Raphael *et al*., 2010). In short, spinal interneurons and synapses of sensorimotor circuits were controlled by simple SET and GO step commands. The SET commands shape the sensorimotor circuits (synaptic gains, level of activity of neurons) without producing any movement, and the GO commands trigger and elicit the movement. Since these signals lacked any dynamic content, the spinal sensorimotor circuit was solely responsible for shaping the dynamics of the commands, particularly in producing the triphasic commands.

We also wanted the neuromechanical system to self-organize its commands as a function of the produced movements, as would do a child exploring the commands of its own arm. Recent advances in developmental machine learning (Baranes & Oudeyer, 2013; Forestier *et al*., 2017) have shown that a novel form of exploration algorithms, called *Goal Exploration Processes* (**GEP**), can be used to automatically define target behaviors and efficiently discover a map of reachable behaviors, thereby solving problems of unreachable behaviors and local minima encountered by alternative, state-of-the-art optimization methods (Oudeyer & Kaplan, 2007; Forestier *et al*., 2017; Colas *et al*., 2018) ; (for related methods using diversity search to solve problems with local minima or sparse rewards, see also Lehman & Stanley, 2011; Mouret & Doncieux, 2012; Pathak *et al*., 2017; Eysenbach *et al*., 2019). Moreover, self-conducted exploration of high-dimensional, continuous parameter systems have the advantage to be sample efficient, i.e., they enable the discovery of diverse behavioral domains with a limited number of experiments. So far, such algorithms have been used to discover diverse behaviors in robotics (Forestier *et al*., 2017; Péré *et al*., 2018) as well as in complex systems such as cellular automata, morphogenetic models (Reinke *et al*., 2019; Etcheverry *et al*., 2020) and chemical systems (Grizou *et al*., 2020), implementing a form of discovery assistant that can leverage interactively guidance of users (Etcheverry *et al*., 2020). Furthermore, GEP is not only an exceptional tool for exploring behavioral domains, but this type of exploration, where an agent defines its own goals, has recently been recognized as a curiosity-driven learning mechanism. This approach is used in artificial intelligence and machine learning, similar to how it occurs in animals and humans (Gottlieb & Oudeyer, 2018).

Finding new sets of parameters and behaviors from previously achieved behaviors has already been successively used to generate intermediate movement using interpolation (Tsianos *et al*., 2014). However, due to the large redundancy of the neural circuits, interpolation fails if previous behaviors relate to different strategies (Tsianos *et al*., 2014). To get a systematic exploration of the capacities of the neuromechanical model, we used a simple GEP protocol inspired from Forestier et al. (2017). These algorithms use two work spaces: (i) one for the behaviors: here we used a two dimensions space (movement amplitude and movement speed), and (ii) one for the parameters that determine the motor commands from the brain: here we used a series of step commands that modify synaptic strength and neuron activity in spinal cord circuits either in the preparatory phase (SET) or to launch the movement). The GEP finds new valid behaviors by repeating 4 stages: (1) Choose a goal (a goal is a targeted behavior); (2) Find the parameters that produced a behavior close to the aim in the data base of preceding successes and modify slightly these parameters in a random way; (3) Run the model with these parameters; (4) Observe the result and if acceptable, store behavior and parameters in the database.

Several modelling approaches have proposed that the CNS employs a minimum jerk strategy when planning any given movement (Flash & Hogan, 1985). This proposal was refined later (Nakano *et al*., 1999) showing that minimum angle jerk predicts the actual arm trajectory curvature better than the minimum jerk model. Therefore, we used it here. Once valid behaviors were found, the GEP searched new ones by modifying preexisting valid behaviors. Because the policy was random, we called this process rGEP. Here, rGEP proved efficient at discovering step commands that enabled spinal circuits to handle a broad spectrum of functional behaviors while eliciting natural triphasic commands over the whole range of the behavioral domain.

## Methods

### A. Software’s used in simulations

During this work, we have used several models of sensorimotor systems controlling the same musculo-skeletal apparatus. All models were developed with the **AnimatLab software** (https://animatlab.com) (Cofer *et al*., 2010), a neuromechanical environment based on user-friendly graphic interface that can be downloaded from https://www.animatlab.com. Execution of AnimatLab models can be controlled by **Python scripts**, which were used to command execution of model movements by **step command** targeting spinal neurons and synapses (see below). All scripts used in the present work are written in Python3.8 language. For a quick overview of GEP principles, see pseudo script in Supplementary materials. Python scripts for GEP and all simulations included in this study can be downloaded from GitHub, with an installation procedure for all software package involved: (https://github.com/Cattaert/rGEP/tree/main).

### B. Musculoskeletal system

The musculoskeletal model consisted of an upper limb involving an elbow actuated by two antagonistic muscles (biceps and triceps) (Figure 1A). This single musculoskeletal model was used in all simulations, with various spinal sensorimotor circuits and step commands controlling their state (see below). Information about this musculoskeletal model is given in table 1.

**Figure 1:**
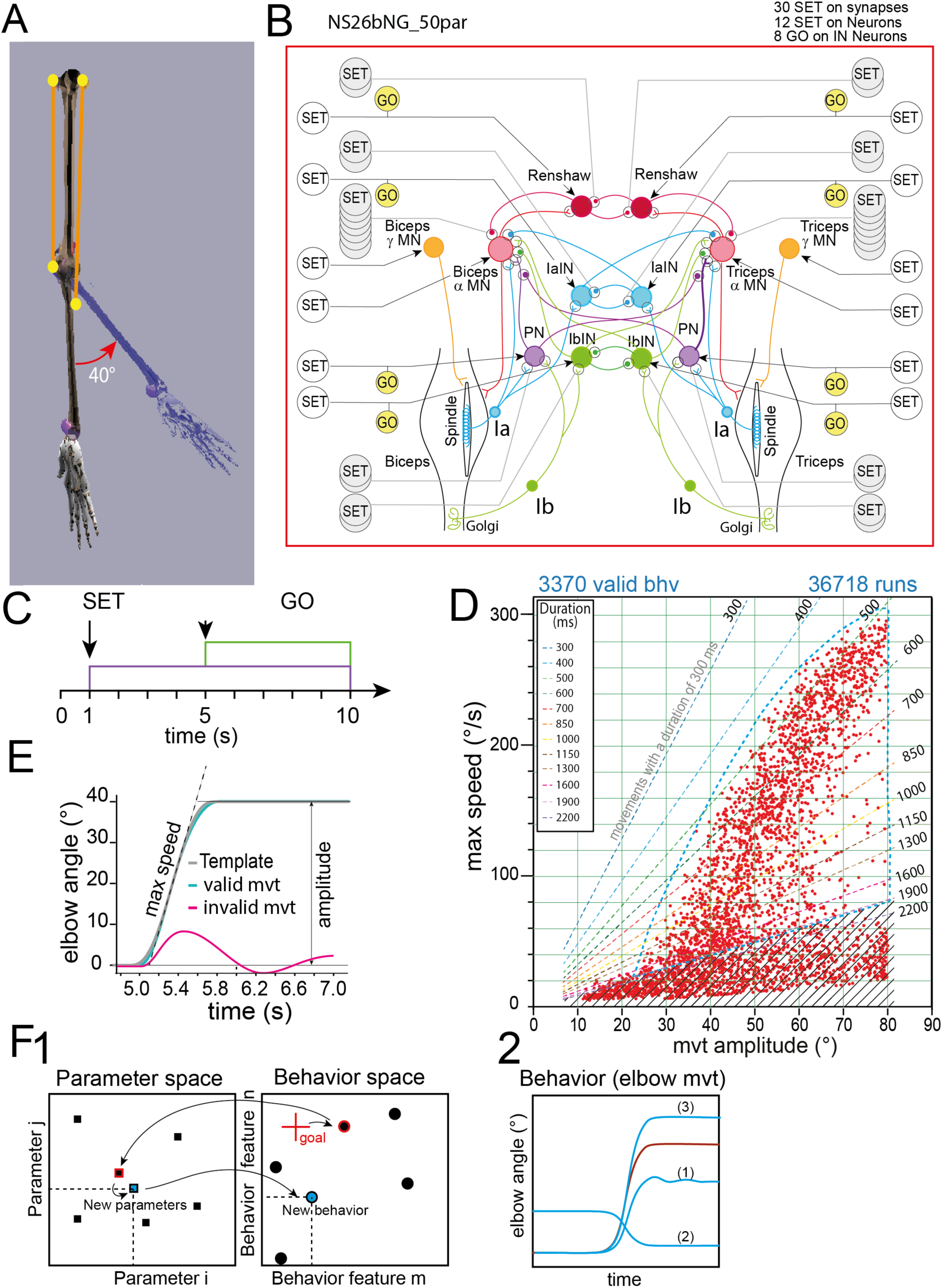
Neuromechanical model and GEP principles. ***A***, Musculoskeletal system. ***B,*** Sensorimotor system comprising αMNs, γMNs, propriospinal interneurons and Ia afferents from muscle spindles, with SET ad GO commands. In this study, simplified sensorimotor networks have been associated to the same musculo-skeletal system (see Figs 4-8). ***C,*** Timing of SET and GO commands used to control the sensorimotor network. The SET commands adjust all neuron activity levels at 1 sec to bring the network in a state ready to trigger the movement (without producing the movement by itself). The GO commands launch the movement. SET and GO commands are the parameters of **parameter space**. ***D***, Example of Behavior space obtained with rGEP algorithm after 39160 iterations. Valid behaviors are indicated by red dots, and corresponding movement durations are indicated by doted lines. ***E***, In this work the **behavior space** was defined by the amplitude of elbow movement (abscissa) and its maximal speed (ordinate). Only, movement (green curve) fitting to a minimum jerk profile (gray curve) were accepted (i.e., mean square error (MSE) < 1 - see Methods). An example of rejected movement is presented (red curve). ***F***, Schematic diagram of the GEP exploration algorithm. **F1**: Process starts from an existing **parameter space** (black filled squares in *Parameter space* – for simplicity reduced to two parameters), which is associated with a **behavior space** (black filled circles in *Behavior space*), a new *goal* is given in the periphery of the known behavioral space (red cross). The algorithm searches for the closest behavior (closest neighbor), it will retrieve the parameters associated with this behavior and replay its parameters including additional noise (“*New parameters”*). This new set of parameters generates a “*New behavior”* in the space of behaviors that is kept or not, depending on its *Cost* value). **F2**: The movement that was closest to the goal (red circle in behavior space, and red movement trace in F2) is a valid movement. However, when its parameters are changed randomly different issues are possible: i) the obtained new movement is invalid, either because there are oscillations (1) or because it is performed in the wrong direction (2); ii) the new movement is valid and may be close or far from its parent.

**Table 1:**
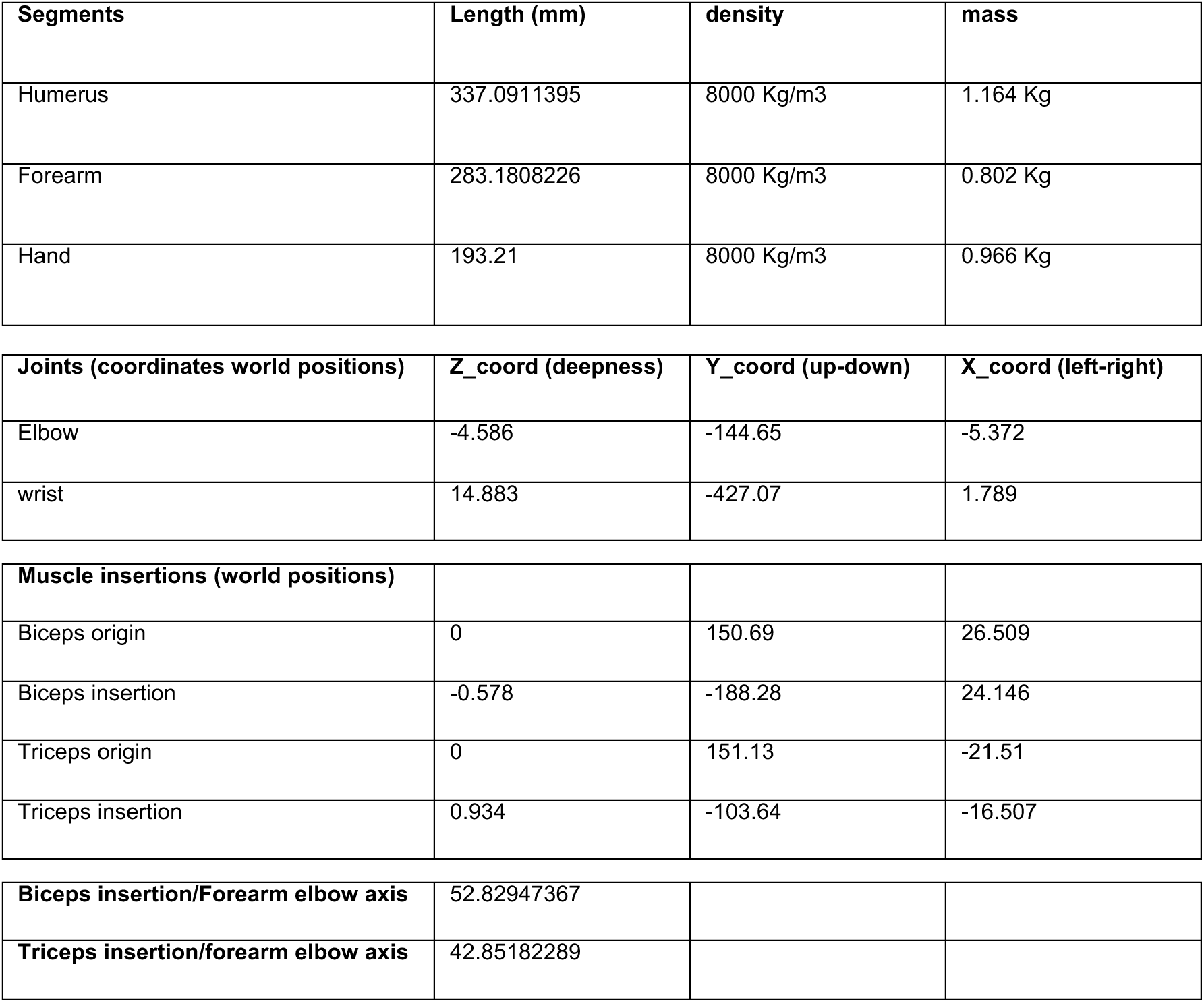
musculo-skeletal model data. Coordinates and dimensions are given in mm. The origin of coordinates (0,0,0) is the center of the Humerus, that is in vertical position at rest (see Fig. 1A).

All movements were produced **in the absence of gravity** in order to avoid natural slowdown of movement speed as the forearm is flexed. Muscles used in AnimatLab are based on the Hill muscle model (Scovil & Ronsky, 2006). The parameters of the Hill model for biceps and triceps are given in Table 2 with length/tension and stimulus/tension parameters.

**Table 2:** Hill muscle parameters used in models for Biceps and Triceps.

|  |  | Biceps | Triceps |
| --- | --- | --- | --- |
| <u>B</u> |  | <u>1 Ns/m</u> | <u>1 Ns/m</u> |
| <u>Kpe</u> |  | <u>100 N/m</u> | <u>100 N/m</u> |
| <u>Kse</u> |  | <u>1000 N/m</u> | <u>1000 N/m</u> |
| <u>Length</u> |  | <u>33.898 cm</u> | <u>27.25 cm</u> |
| <u>MaxTension</u> |  | <u>400 N</u> | <u>300 N</u> |
| <u>Length-Tension</u> | <u>Lwidth</u> | <u>20 cm</u> | <u>11 cm</u> |
|  | <u>Resting Length</u> | <u>40 cm</u> | <u>38 cm</u> |
|  | <u>Range of length</u> | <u>27 – 34 cm (0-100°)</u> | <u>27.5– 32.5 cm (100-0°)</u> |
|  | <u>Range of Tension%</u> | <u>58% - 92%</u> | <u>2% - 70.5%</u> |
| <u>Stimulus-Tension</u> | <u>Amplitude</u> | <u>400 N</u> | <u>300 N</u> |
|  | <u>Steepness</u> | <u>100</u> | <u>100</u> |
|  | <u>X offset</u> | <u>-20 mV</u> | <u>-30 mV</u> |
|  | <u>Y offset</u> | <u>-4.5 N</u> | <u>-4 N</u> |

### C. Presentation of the sensorimotor networks and their control by step commands

Non-spiking (NS) neurons were used to simulate neurons in spinal sensorimotor networks. NS neurons allow a continuous synaptic activation that depends only on presynaptic threshold potential, presynaptic saturation potential, equilibrium potential for post-synaptic effect and synaptic gain (maximal synaptic conductance). The change in postsynaptic conductance increases from 0 (at presynaptic threshold potential) to the maximal synaptic conductance (for presynaptic saturating potential and over). Between these presynaptic threshold and saturation potentials, the postsynaptic conductance increases linearly. Such neurons present the advantage to avoid discontinuity of spikes and spiking synaptic transmission. This is specially critic in small neuron networks, when the level of activity is low. Using NS neurons can also be considered as representing population neurons instead of single neurons.

In this study, we have used several sensorimotor spinal networks, all derived from the “complete” model (Figure 1B). The complete spinal network (50 parameters model, Figure 1B) received proprioceptive feedback from muscle spindles (Ia) and Golgi Tendon Organ (GTO). The associated sensorimotor circuits were inspired from Raphael et al (2010), and contained 12 neurons (2 alpha motoneurons (MNs), 2 gamma MNs, 2 propriospinal interneurons (INs), 2 Ia INs, 2 Ib INs, and 2 Renshaw INs), and 30 synapses.

The specific aim here was to generate minimum jerk movements (i.e., fitting to this movement shape was in the cost function – see below). We used the same principles for step commands as Raphael *et al*. (2010). Step commands were represented as step functions starting at t=1s (SET command) and at t=5s (GO command) (Figure 1C). The SET commands were responsible for the state of readiness of the network. This first stimulation should activate the network without releasing any movement (Prut & Fetz, 1999; Eliasmith & Anderson, 2003).

The SET command targeted all the elements of the network (neurons and synapses) and their control was maintained constant during all the time of the simulation (up to t=10s). The GO commands were responsible for movement initiation and were maintained constant until the end of the simulation. In this example GO commands modulated the level of activity of propriospinal interneurons (PN), 1a interneurons (IaINs), 1b interneurons (IbINs) and Renshaw INs (Renshaw) (see Figure 1B). Note that at t = 5 seconds, the transient changes induced by the SET command had stabilized before the GO command was issued. In summary, GO commands targeted the 8 INs, SET commands targeted the 12 neurons and adjusted the 30 synapses’ gains.

### D. Brief overview of rGEP

To assess behavior capabilities associated with a given neuromechanical model (i.e., the variety of movements that can be produced (Figure 1D) and meeting some criteria like minimum jerk - see below for the **cost function**), we have developed a rGEP algorithm that uses two spaces (the parameter space and the related behavior space). The movements contained in the behavior space are defined by their amplitude and their maximal speed (Figure 1E). At each iteration, goals (i.e., movements of given amplitude and speed) are self-produced by the algorithm. Goals are disposed either on the periphery of (extend strategy) or inside (fill strategy) the known behavior space. In the illustration of Figure 1F1, the example goal is on the periphery of the behavior domain (see red cross). The closest behavior is then searched and its associated parameters are slightly modified (**random function**) to produce a new parameter set (*New parameters*). We choose a random policy in order not to introduce bias in the setting of new values for parameters. Because the policy used by GEP for parameter changes was a random function, this GEP was named **rGEP.** After a simulation is run (Figure 1F2), if the produced movement fulfils the **acceptance criteria**, it is stored in the behavior domain (see trace (3) in Figure 1F2), otherwise, it is rejected (traces (1) and (2)). Acceptance criteria were as follows: (i) the produced movement should fit a **minimum jerk angular kinetics** (Friedman & Flash, 2009); (ii) there should be no oscillations in preparatory and post movement phases; (iii) the movement amplitude should be in the range (10-110°); (iv) no co-contraction should be present during the preparatory phase (controlled by SET commands, see below) and during the stabilized phase after movement (launched by GO commands). A **cost function** ensured valid behaviors fulfill acceptance criteria (see methods)

Because rGEP needs valid behaviors to progress, the first step, was to find initial valid behaviors. In order not to interfere with the design of the commands, these initial behaviors (termed **seeds**) were found via a state of the art optimization method (covariance matrix association evolution strategy (CMAes) (Hansen *et al*., 2003). Briefly, a desired behavior (a movement template) was given to the optimization method that worked on the parameters of the SET and GO commands to find the result that best fitted the desired movement template (see next paragraph for more explanations). During this process of optimization, any valid movement produced was stored and could be used as a seed.

### E. Optimization method used to get first behaviors (seeds)

Covariance Matrix Adaptation evolution strategy (CMAes) is an evolution algorithm developed by Hansen (Hansen *et al*., 2003). It works with a gradient descent with the aim to minimize a cost function. This evolution algorithm is based on the principle of biological evolution. This means that for each generation (iteration), new individuals are generated from variation, in a stochastic way, of the current generation. To initiate the next generation, individuals with the best scores (minimal value of the cost function) are selected to become the parents of the next generation. Following this rule, individuals of the successive generations get closer to the best results.

In the evolution algorithm, the solutions that represent the new generations are issued from multivariate-normal distribution based on a variance value (*sigma*) and the covariance matrix of the preceding best solutions. This allows the stochastic changes made to the parameters to be oriented in the direction of the solution. Moreover, from one generation to the next, the *sigma* value is adapted: increased when two generations are in the same direction, and decreased if not. This allows accelerating the progress of optimization.

In the CMAes optimization procedure, some parameters have to be given before the simulation was launched including the *limits* of exploration and the *sigma* value. If the *limits* of exploration were too large, new individuals could be generated too far from each other and the optimal solution might not be reached. A small *sigma* will create a new generation not so different from the previous one and a large one will bring more changes. All optimization series were initiated with a small *sigma* (0.005) and *limits* of exploration covering 100% (i.e., between 0 and 1 for all normalized parameter values). *Sigma* and *limits* were given to CMAes at initiation of optimization process.

### F. Cost function used to select valid behaviors

For each movement produced, a **cost function** estimated the proximity of movements produced by the simulation to human movement. As normal human movements follow a template based on the minimum jerk model (Flash & Hogan, 1985), the **cost function** calculates the distance (mean square error) between the produced movement and this template (Figure 1E), with the addition of the penalty for the co-contraction (*Equation 1*). This term was added after a first exploration of the method, which led to systematic co-contraction during the preparation period (result of the SET command). The result was considered valid when the *Loss* value was less than 1.

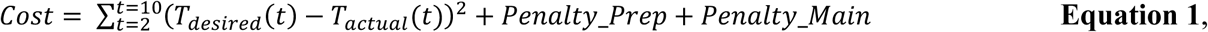

with t representing the different sampled times of the simulation (dt=10ms).

In order to avoid solutions in which co-contraction occurred during the preparatory phase (or after movement stabilization), co-contraction penalty was added in each of these phases. *Penalty_Prep* was calculated during the preparatory phase (start=2s, end=5s), and *Penalty_Main* during the maintained position phase (start=7s, end=10s). The movement phase was not included because complex interactions may occur due to the dynamics of sensorimotor interactions. The penalty for co-contraction was calculated as indicated in *Equation 2*.

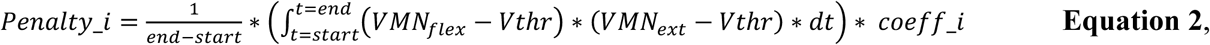

in which *VMN_flex_* and *VMN_ext_* are the membrane potential of the flexor and extensor MNs, respectively, and *V_thr_*=-60mV, is a threshold voltage under which penalty = 0.

The coefficient (*coeff_i*) was set to 100 during the preparatory phase to make sure coactivation would be rejected in this phase, and *coeff_i* was set to 0 during the maintained position after movement to allow coactivation in this last phase.

### G. Time course of the rGEP algorithm

An example of rGEP time course is presented in Figure 2, on the same model presented in Figure 1 (see also inset network diagram in Figure 2). Before rGEP starts, **valid behaviors (seeds)** must be given to the algorithm (see below). In the example presented in figure 2, four seeds were given (see blue crosses in Figure 2A1-4).

**Figure 2:**
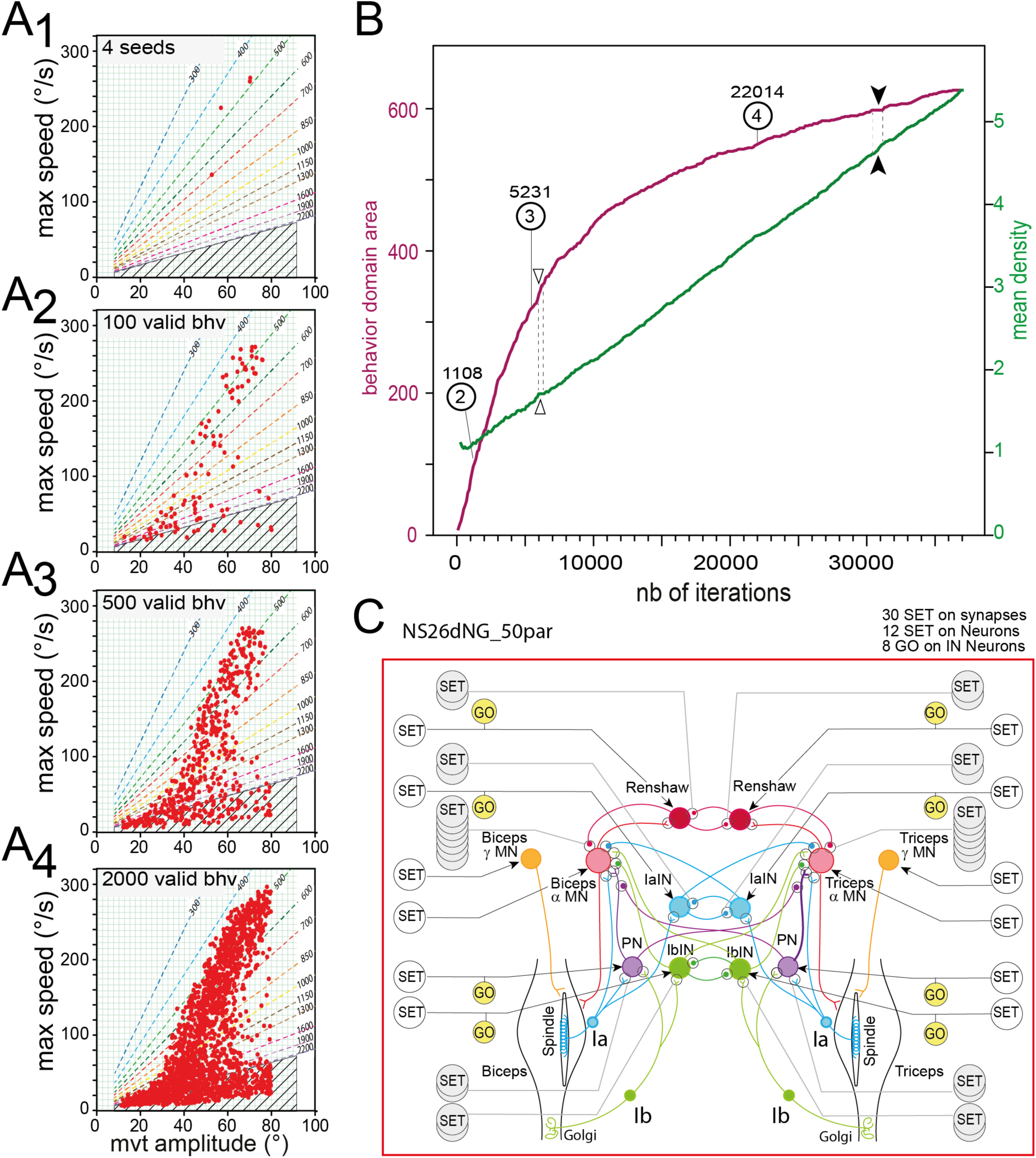
Temporal evolution of rGEP. **A**, Progression of the behavior domain uncovered by rGEP after starting with 4 seeds (A1), and extending to 100 (A2), 500 (A3) and 2000 (A4) valid behaviors. **B**, Time course of behavior domain area expansion (i.e., number of squares containing at least one valid behavior in the green grids presented in A1-4) presented in violet trace. The mean density of behaviors in the behavior domain area is presented in green trace. The numbers 2, 3 and 4 correspond to the behavior domains representations shown in (A2-3). **C**, Diagram of the 50-parameter model.

Four pictures of the behavior domain extension are presented starting from 4 seeds (Figure 2A1), and extending to 100, 500 and 2000 valid behaviors (Figure 2A2-4), corresponding to 394, 1924 and 6926 runs of simulation, respectively. As the rGEP progresses, the behavior domain extends (Figure 2B, violet trace, *behavior domain area*). The behavior domain area was quantified as the number of cases of a grid, defined in the behavior work space (see Methods), which contain at least one valid behavior (see green grids in Figure 2A1-4).

While the borders of the behavior domain get more and more precise, there were still some empty central zones when the process was stopped after 300 “extend” rounds and 50 “fill” rounds (totalizing 36718 runs). Note that there is no direct relationship between the number of extend rounds and the number of runs, because the number of runs in a given round depends on the size of the behavioral domain at the time the round is launched. Note also that when the rGEP was stopped, the process was almost complete as indicated by the very slow increase of behavioral domain area (violet trace) during the last 15000 runs (Figure 2B). The behavior domain at the end of the 300 extend and 50 fill rounds (36718 runs) was presented in Figure 1B. We note that some empty zones still persist after the 3000 extend rounds. This domain was then completed by adding three new rGEP with 300 expands and 50 fills (starting from new seeds each time) totalizing 112240 runs (see below “*Behavior domain of the complete (50-parameters) model*”).

We can also see that the progression of rGEP is not linear. Indeed, several transient plateaus were observed (see filled arrow heads in Figure 2B) on the behavior domain area curve (violet trace). During such blockage of exploration, valid results were found in the already discovered behavior domain, resulting in a transient increase of the mean density (see methods) of valid behaviors (green curve in Figure 2B). The situation unblocks when an original valid behavior is found that extends the behavior domain, and allows new behaviors to be found from its parameter set, resulting sometimes in an abrupt increase of the behavior domain area (see open arrow heads in Figure 2B) while the progression of mean density slows down (see green trace, in Figure 2B). As the behavior domain extends (Figure 2A-B), the parameter domain also evolves.

The final parameter domain is presented in Figure 3. In this figure, all valid values of the 50 parameters are presented in 25 graphs, each displaying a pair of parameters: 4 GO stim pairs: [(*Triceps PN GO, Biceps PN GO), (Triceps 1aIN GO, Biceps 1aIN GO), (Triceps 1bIN GO, Biceps 1bIN GO), (Triceps Renshaw GO, Biceps Renshaw GO)*] (Figure 3A1-4); 6 SET stim pairs: [(*Triceps Alpha SET, Biceps Alpha SET),* (*Triceps PN SET, Biceps PN SET), (Triceps 1aIN SET, Biceps 1aIN SET), (Triceps 1bIN SET, Biceps 1bIN SET), (Triceps Renshaw SET, Biceps Renshaw SET), Triceps gamma SET, Biceps gamma SET*] (Figure 3B1-6); 15 SET syn pairs: [*Ext 1a-> ExtAlpha*, *Flx 1a->FlxAlpha*] (Figure 3C1), [*Ext 1a->ExtPN, Flx 1a>FlxPN*] (Figure 3C2), [*Ext 1b- >PN, Flx 1b- >PN*] (Figure 3C3), [*Ext PN- >ExtAlpha, Flx PN- >FlxAlpha*] (Figure 3C4), [*Ext 1a- >Ext1aIN, Flx 1a- >Flx1aIN*] (Figure 3C5), [*ExtAlpha->ExtRenshaw, FlxAlpha->FlxRenshaw*] (Figure 3C6), [*ExtRenshaw->ExtAlpha, FlxRenshaw->FlxAlpha*] (Figure 3C7), [*ExtRenshaw-> FlxRenshaw, FlxRenshaw-> ExtRenshaw*] (Figure 3C8), [*Ext1bIN- > FlxAlpha, Flx1bIN- > ExtAlpha*] (Figure 3C9), [*Ext1b- >Ext1bIN, Flx1b- > Flx1bIN*] (Figure 3C10), [*Ext1bIN- >Flx1bIN, Flx1bIN- > Ext1bIN*] (Figure 3C11), [*Ext1bIN- >ExtAlpha, Flx1bIN- > FlxAlpha*] (Figure 3C12), [*Ext1bIN- >FlxAlpha, Flx1bIN- > ExtAlpha*] (Figure 3C13), [*Ext1aIN- >Flx1aIN, Flx1aIN- > Ext1aIN*] (Figure 3C14), [*ExtPN- >FlxPN, FlxPN- > ExtPN*] (Figure 3C15), in which *stim* = stimulus sent to a neuron, and *syn* = strength setting of a synapse between a presynaptic and a post synaptic neuron separated by “->” (see Figure 3 legend for more explanations).

**Figure 3:**
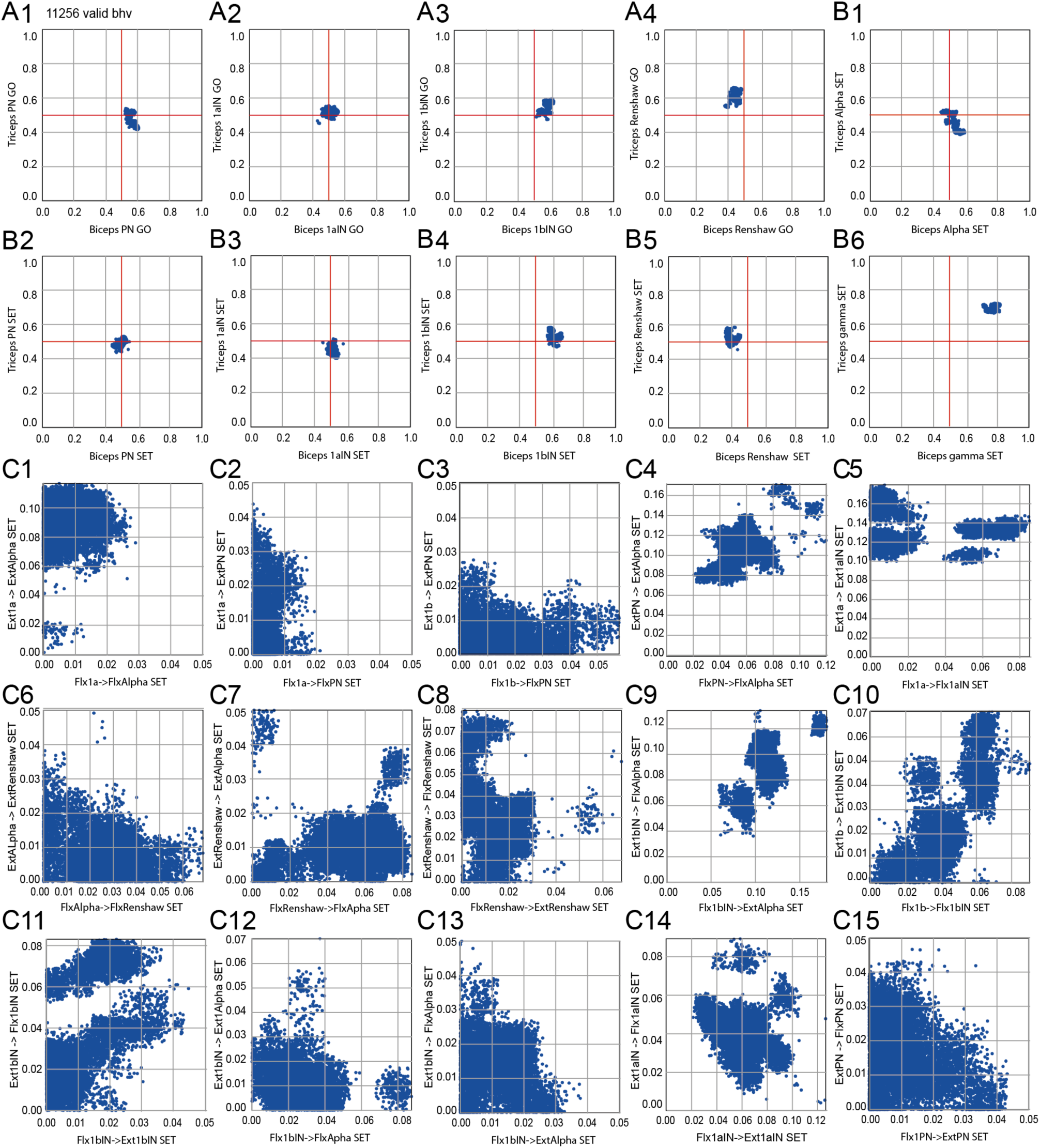
Parameter domains of the complete model. The 50 parameters are represented in 25 graphs displaying 2 parameters each: 4 stimulus GO graphs (A1-4), 6 stimulus SET graphs (B1-6) and 15 synapse strength graphs (C1-15) obtained after 39160 runs. Axis names: stimulus graphs represent intensity of stimulus sent to a neuron in the GO (A1-4) and SET (B1-6) commands; synapse strength graphs represent strength setting of a synapse between a presynaptic and a post synaptic neuron separated by “->” (for example Flx1a->FlxAlpha SET indicate the SET command for synaptic strength between 1a spindle afferent and alpha MN).

### H. Modelling coding properties of muscle spindle

Muscle spindles were added to each muscle. In AnimatLab (see Methods), muscle spindles have the same “attachments” as the muscles. Their tension-length ratio, maximum force capacity, and viscoelasticity are based on the Hill muscle model (Scovil & Ronsky, 2006). To assess the quality of this neuro-musculoskeletal model, imposed movements of flexion and extension were simulated, while muscle spindle activity was recorded (Figure 4A). Our first work was to adjust by hand the muscle spindle parameters that controlled their coding of movements so that their response were comparable to electrophysiological data from the literature (Boyd *et al*., 1977) (Figure 4B), in the absence and in the presence of activation of a Gamma MN.

**Figure 4.**
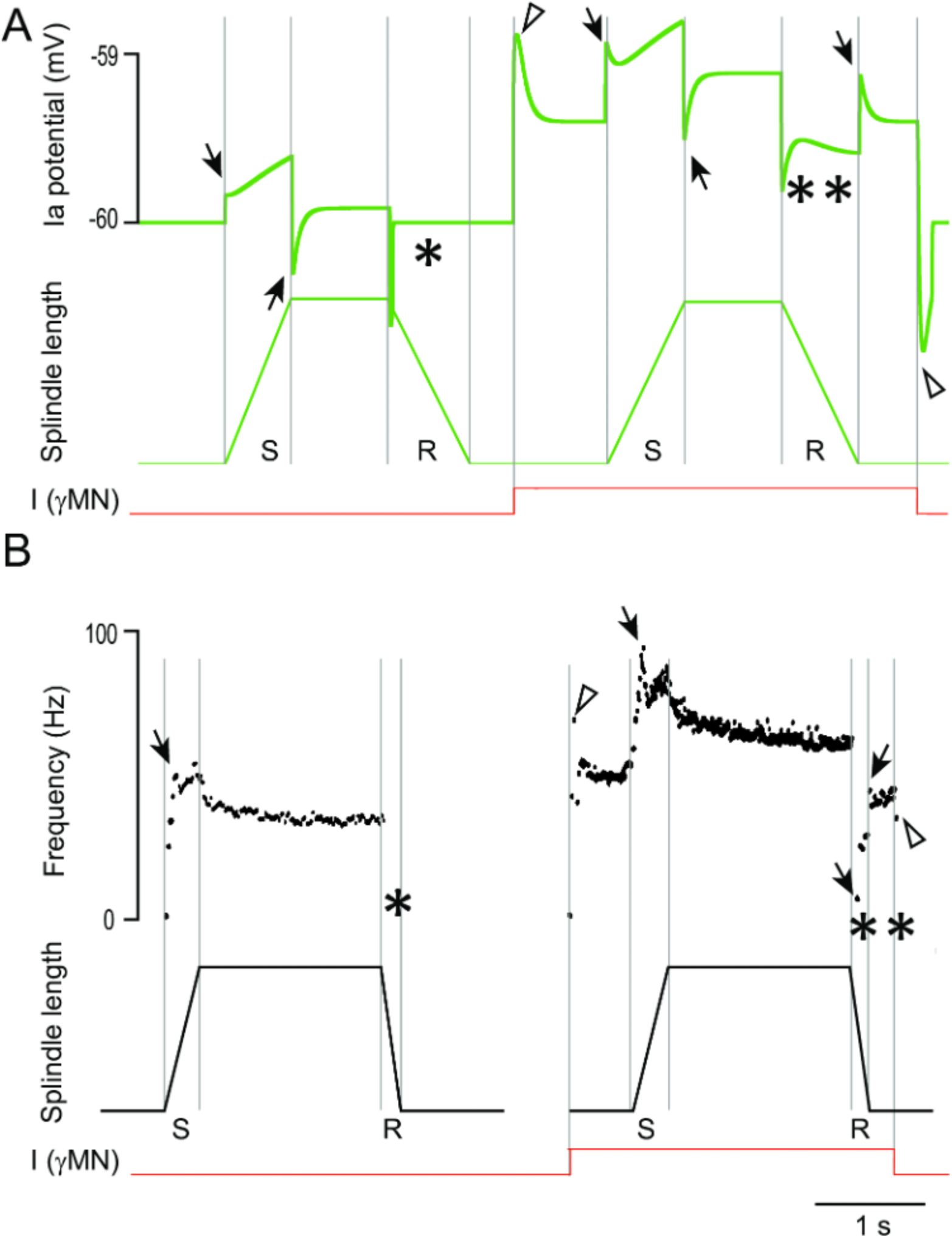
Muscle spindle coding properties. **A.** Modeled extensor muscle spindle activity during stretch (S) and release (R) ramps, without (left), and with (right) activation of the gamma motoneuron (I γMN). **B.** Electrophysiological recording of the Ia fiber during the same imposed movements as in **A** modified from (Boyd *et al*., 1977) Note that, in both simulation and physiological recording, the first release of the spindle does not give any response (see *) due to lack of tension of the spindle. However, the activation of the γMN allows the spindle to give an answer during the second release (see **). Note also that in **A**, the “acceleration” component produced by the serial spring velocity (in the Hill model) is responsible for the transient activation at each movement change (arrows), while the “tonic” component produced by the tension of the spindle (in the Hill model) produces an exponential decay shape. Transients are also produced at onset and offset of the γMN stimulation. These features are present in **B** except for transients at the end of stretch movements.

Note that the AnimatLab platform proposes a single type of gamma command that is a mixture of gamma static and gamma dynamic properties. Therefore, in all simulations we used this single type of gamma MN (see Discussion F for more details).

## Results

### A. Goal exploration process explores the capacity of spinal sensorimotor circuits and their step commands to produce behaviors

In this section, we provide a brief overview of the neuromechanical models and the process (rGEP) used to explore their behavioral capabilities.

#### 1) Behavior domain of the complete (50-parameters) model with both Ia and Ib afferents and associated network

The behavior domain found for the 50 parameters model by the rGEP is presented in Figure 5B. It was obtained by cumulating 4 rGEP rounds, each composed of 300 “extend” rounds and 100 “fill” rounds, and totalizing 112240 runs, in which 37087 behaviors were valid. The behavior domain is defined by two parameters of the produced movements: maximal speed (ordinate) and movement amplitude (abscissa). A third characteristic of movements (movement duration) is presented on the graphs by oblique lines labelled with the represented duration (in ms). Only physiologically plausible movements have been retained (slow movements the duration of which was > 2s have been rejected – see striped area). Only movements with an amplitude in the range [10°, 110°] have been retained, to avoid too small movements and too large movements that could reach the flexion limits. With both Ia and Ib sensorimotor circuits complete, this model could perform large movement (up to 80°) at a rapid speed (>300°/s) with movement durations < 500ms. Note for each amplitude, a great variety of speeds was found.

**Figure 5:**
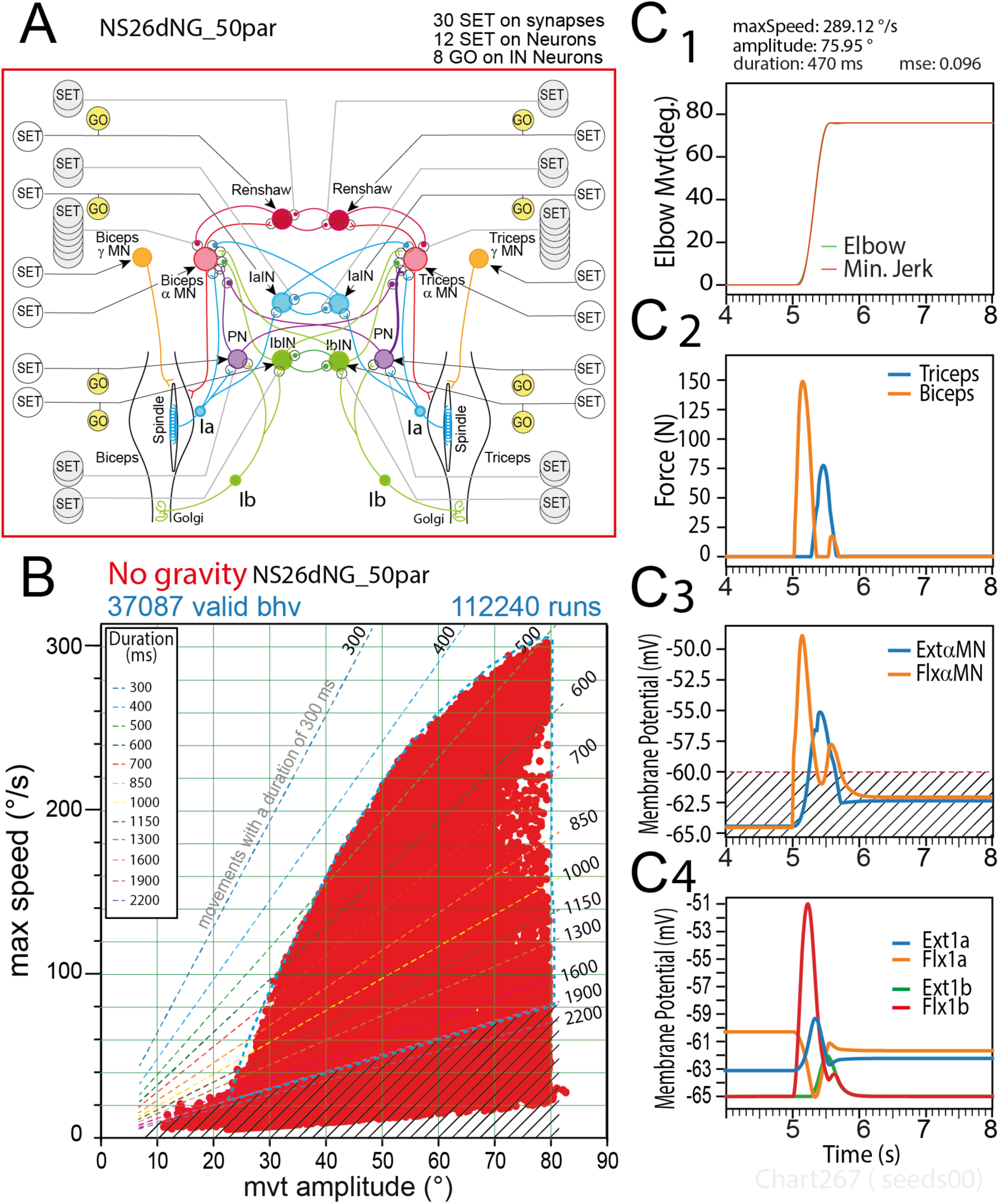
Movements produced by rGEP with the 50-parameters model. **A**, Sensorimotor spinal network of the 50-parameters model. **B**, Behavior domain uncovered by rGEP on this model. The perimeter of the domain is outlined with a blue dashed line. **C**, Representative example of one valid movement in the behavior domain (C1), with force time course of Biceps and Triceps muscles (C2), Flexor and Extensor MN membrane potential (see FlxαMN and ExtαMN in C3), and 1a & 1b afferent neurons potential (see Flx1a, Ext1a, Flx1b, Ext1b in C4). Striped area in C3 indicates MN under-threshold potentials, under which no muscle response was observed.

A representative example of a large movement (amplitude 75.95°) is given in Figure 5C1. Note the profile of the movement that fitted a minimum jerk profile with small MSE value (0.096). Several other variables are also presented for this example: Force, Membrane potentials of MNs, and sensory neurons. At the initiation of the movement, the force recorded in the Biceps was large (reaching >120 N within 110 ms) and decayed to 0 in 200 ms (Figure 5C2). The force recorded in the triceps increased at the end of the rapid decrease of biceps force, and reached 75 N at t=545ms, and then decreased in the next 150 ms, at the end of the flexion movement. Note the small second peak of flexion force while movement stops.

The activity of alpha MNs present transient activities during the movement phase: the biceps MN (*FlxαMN*) presents two peaks of depolarization, the second being smaller, and an absence of activity between the two peaks, during which the triceps MN (*ExtαMN*) presents a peak of activity (Figure 5C3). Once the flexed position is reached, the activity of both alpha MNs totally vanishes. During the movement phase, the proprioceptive feedback from Ib afferents was much larger than that of the Ia afferents (Figure 5C4). The shape of Ib activations was also different from that of Ia afferents: during the rapid flexion movement, the Ib activity of the flexor muscle (*Flx1b*) present a large peak during the acceleration of the flexion movement, in accordance with its role in sustaining the flexion movement via the Flexor PN excitatory circuit (see Figure 5A); this initial *Flx1b* peak is followed by a smaller peak of the extensor 1a activity (*Ext1a*) that contributes to slow down the flexion movement due to its excitatory effect on the extensor αMN (see Figure 5A); this slowing effect is prolonged by the delayed increase of *Ext1b* activity that contributes to the activation of the extensor αMN via the PN interneuron of the extensor circuit (see Figure 5A). Note that during the flexion movement, the activity of the Flx1a presents a trough. Note also the small peak of Flx1a just before stabilization at -62 mV.

The time course of 1a and 1b afferent activities (larger Flx1b peak followed by a smaller Ext1a peak, and by a delayed peak of the Ext1b activity, concomitant with the recovery of Flx1a trough) were regularly observed in the 50-parameter model, with some small variations all along its behavior domain. This first study will serve to compare the capacities of other models issued from this one, but lacking some neural elements. To facilitate comparison, the perimeter of the behavior domain of the 50-parameter model (see dashed blue line in Figure 5B) will be superimposed on their behavior domains.

#### 2) Behavioral domain produced in the absence of Ib feedback (with central cross connections)

In order to study what proprioceptive feedback was necessary to produce valid movements, in a first step, we suppressed the Ib input, the Ib INs and their synapses (Figure 6A). When rGEP was run on this model, the resulting behavior domain was close to the one obtained with the complete spinal network (compare Figure 5B and Figure 6B), although maximal speed for large movements was slightly reduced, and all movement durations were longer than 450 ms.

**Figure 6:**
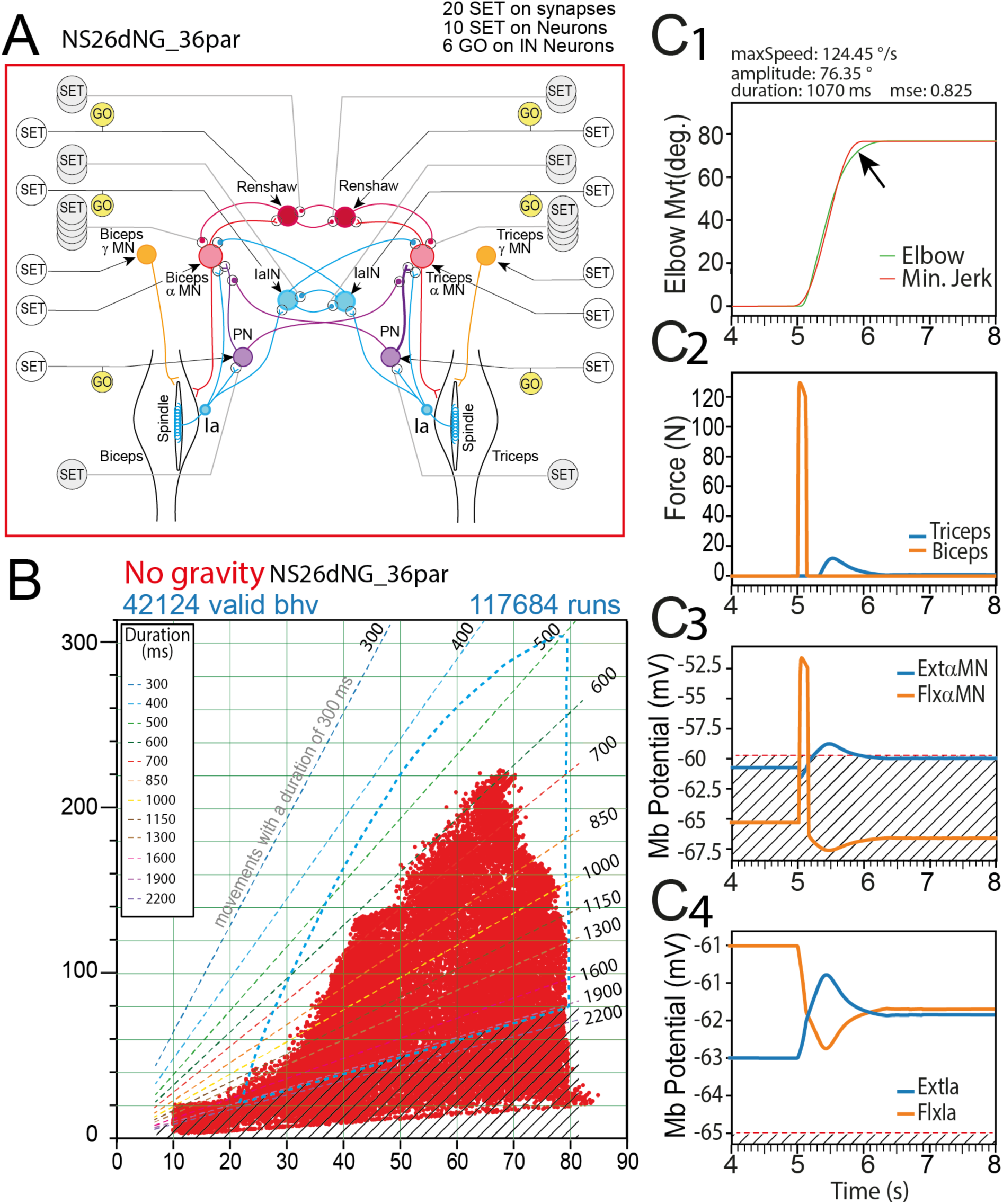
Movements produced by rGEP with the 36 parameters model. **A**, Sensorimotor spinal network of the 36 parameters model. **B**, Behavior domain uncovered by rGEP on this model. For comparison, the perimeter of the domain of the 50-parameter domain is represented with blue dashed line. **C**, Representative example of one valid movement in the behavior domain (C1), with force time course of Biceps and Triceps muscles (C2), Flexor and Extensor MN membrane potential (see FlxαMN and ExtαMN in C3), and 1a afferent neurons potential (see Flx1a, Ext1a in C4). Striped area in C3 indicates the potentials under which no muscle response mas observed. Striped area in C4 indicates under threshold activity.

Forces, MN activities and muscle spindles feedback (Figure 6C) were very similar to that of the complete model (Figure 5C) for comparable movements. A representative example of a large movement (amplitude 76.35°) is given in Figure 6C1, with a maximal speed of 124.45°/s. However, the fitting to a minimum jerk template (MSE=0.825) was not as good as in the complete model. This was likely due to the slowdown at the end of movement (see arrow in Figure 6C1) being much more progressive than in the complete model. The amount of force developed by the Biceps muscle (Figure 6C2) was slightly smaller (#130 N) than that observed for the complete model. Strikingly, in this model deprived of Ib afferent circuits, the second peak of Flx muscle force and Flx Alpha MN activity was never observed (Figure 6C2-3). This second peak was also absent from the Flx1a activities (Figure 6C4).

#### 3) Behavioral domain produced in the absence of Ib feedback (without central cross connections)

To go further in the analysis of the involvement of Ia proprioceptive feedback in the production of minimum jerk movements, we drastically reduced the sensorimotor network, by suppressing all relationships between extensor and flexor parts of the network, as well as all interneurons except propriospinal INs. The resulting spinal network is the 14-parameters simplified network presented in Figure 7A with the same design as other models presented in Result section. Interestingly, this extremely simplified spinal network, reduced to Ia proprioceptive inputs connecting their homonymous MN either directly or via their propriospinal IN (PN), was capable of producing some valid behaviors (Figure 7B). The resulting behavior domain was, however much smaller than the one of the 36-parameters mode (no movement shorter than 1150 ms were produced and maximal speed was under #100°/s). A representative behavioral example (amplitude 84.29°) is given in Figure 7C1. Note that in most of the behaviors produced by this model, the fitting to a minimum jerk template was not as good as in the complete model (here MSE=0.725). Moreover, the second peak in Flx force, Flx alpha MN and Flx1a activities was never observed (Figure 7C2-3).

**Figure 7:**
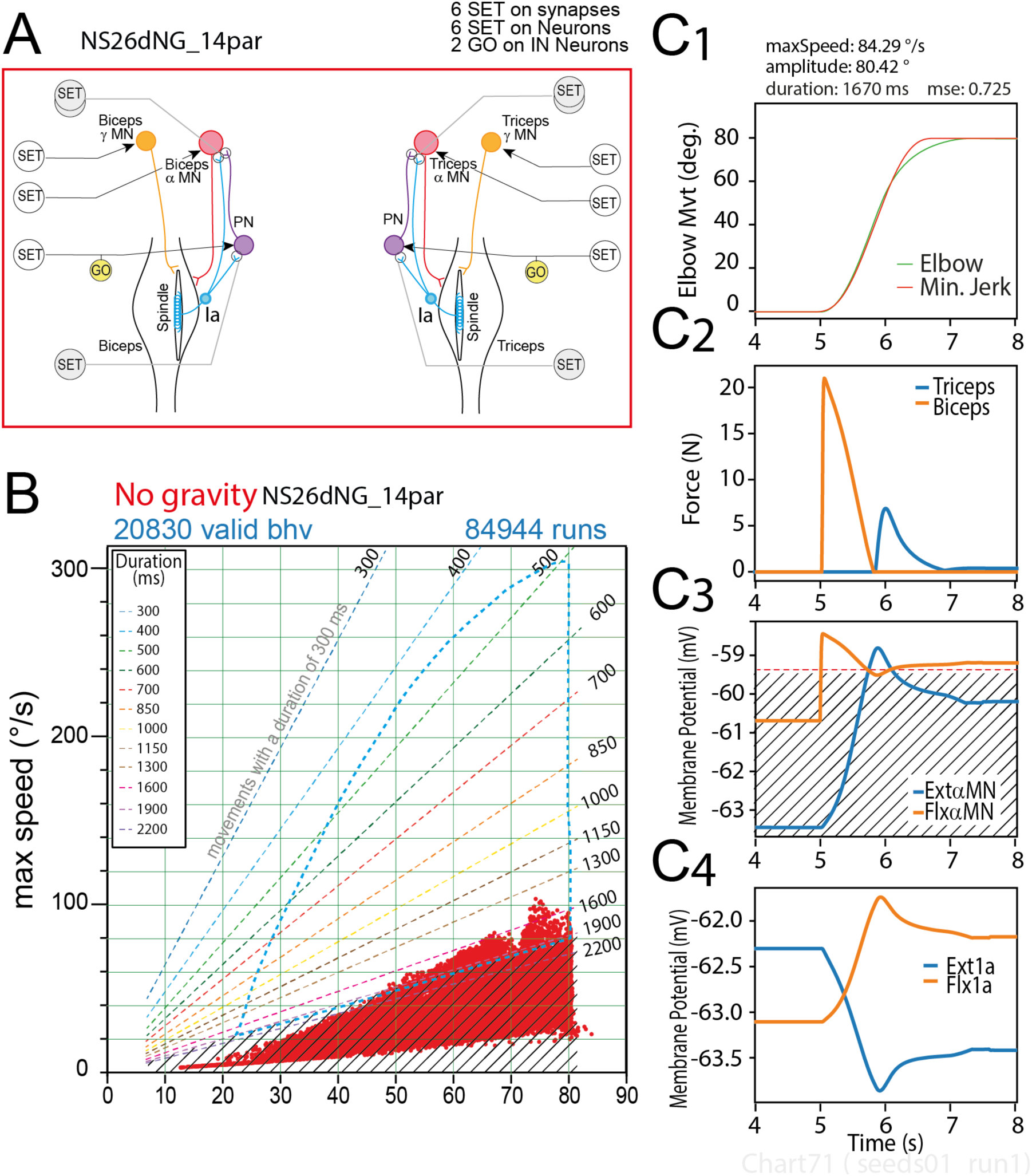
Movements produced by rGEP with the 14-parameters model. **A**, Sensorimotor spinal network of the 14 parameters model. **B**, Behavior domain uncovered by rGEP on this model. For comparison, the perimeter of the domain of the 50-parameter domain is represented with blue dashed line. **C**, Representative example of one valid movement in the behavior domain. Striped area in C3 indicates the potentials under which no muscle response mas observed.

#### 4) Behavioral domain produced in the absence of Ia feedback with central cross-connections

We then applied the same method to test Ib afferents involvement in the production of valid movements via rGEP. Starting from the complete (50-parameters model), we suppressed Ia afferents, Ia INs and their synapses (Figure 8A). This Ib network could not produce any valid behaviors (Figure 8B).

**Figure 8:**
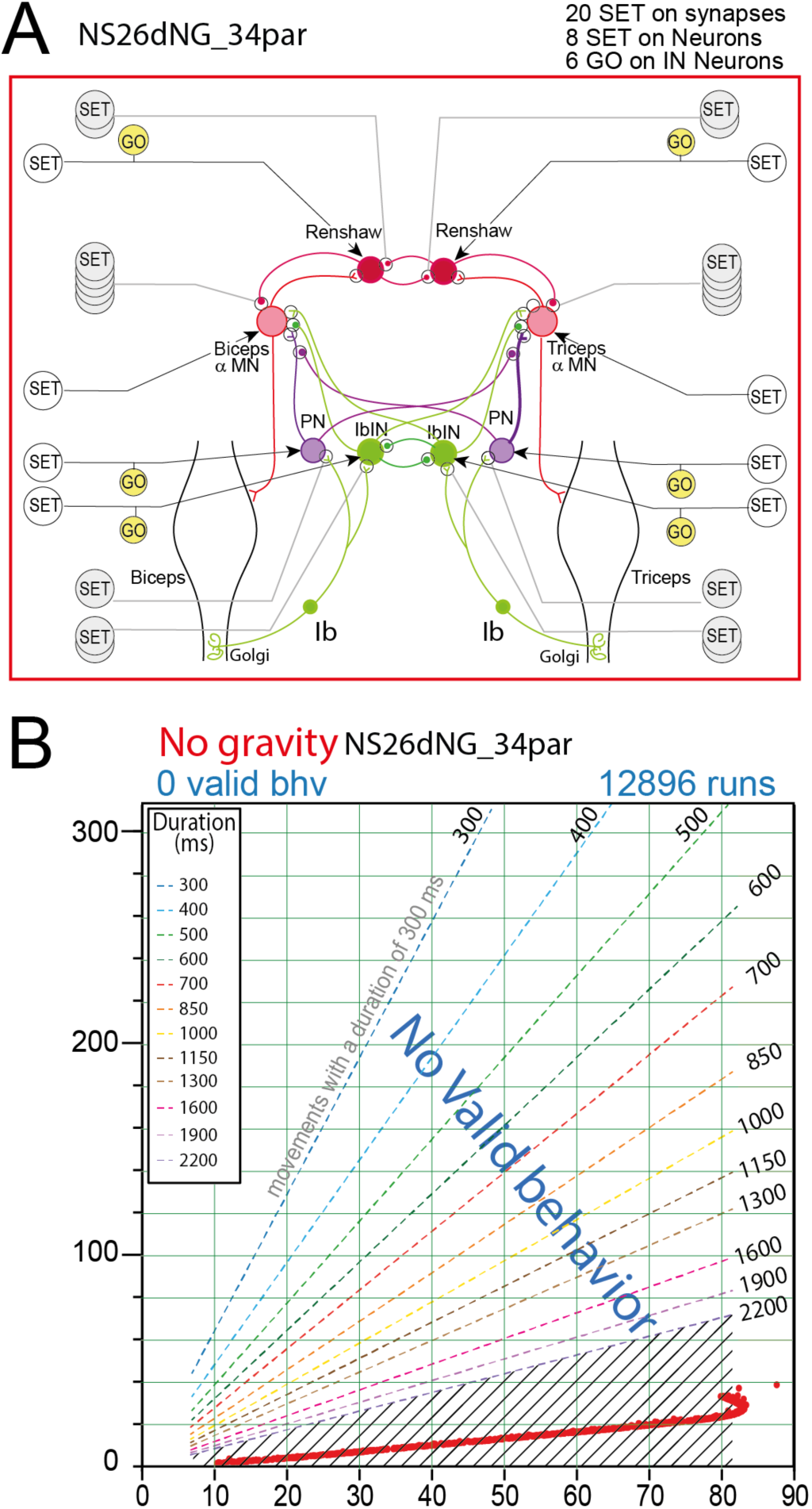
Movements produced by rGEP with the 34-parameters model. **A**, Sensorimotor spinal network of the 34 parameters model. **B**, Behavior domain uncovered by rGEP on this model. The very slow movements produced by this model did not meet criteria for valid movements.

#### 5) Behavioral domain produced in the absence of any feedback but with all INs

Because reciprocal connections seemed important to perform valid behaviors, we tested if complete spinal network with all reciprocal connections but deprived of any proprioceptive feedback (Figure 9A) could produce valid behaviors. However, no valid behaviors were found with this model (Figure 9B).

**Figure 9:**
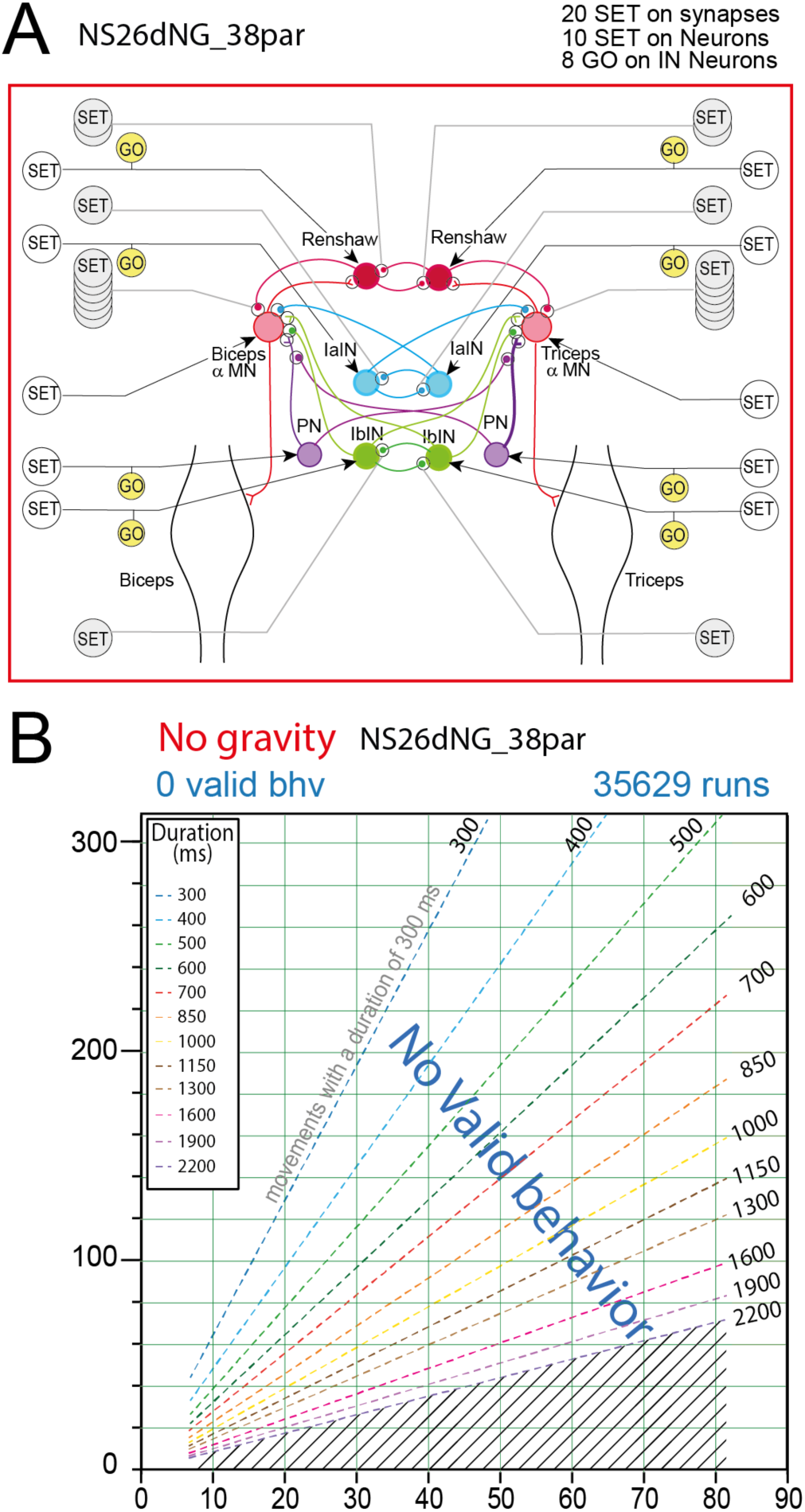
Absence of Valid Movements produced by rGEP with a complete model deprived of proprioceptive feedback. **A**, Sensorimotor spinal network of the 38-parameters model. **B**, Absence of any valid behavior in the behavior domain uncovered by rGEP on this model

### B. Analysis of triphasic patterns in the various models

By design, valid behaviors found by rGEP followed the minimum jerk rule. However, as we will see, this rule does not imply conformity with a triphasic command. Triphasic command is defined from the EMG recordings of two antagonistic muscles that were involved in a rapid movement around a joint (Wachholder, 1928; Angel, 1974; Wadman, 1979; Brown & Cooke, 1981). Typically, the agonist muscle produced two bursts of activity: one (AG1) at the onset of movement and one (AG2) at end of the movement. The amplitude of the initial agonist burst (AG1) was larger than that of the second (AG2) (Bullock, 1992). In between the two agonist bursts, an antagonist burst (ANT) is present (Sanes & Jennings, 1984), the role of which is to slow down the movement speed so that, ideally, the speed is 0 when target angle is reached. However, in rapid movements, if the activity of the antagonistic muscle is too large, the movement goes back. The asymmetrical rise- and fall-times inherent to muscle activation may also induce a prolonged effect of ANT. These effects are counteracted by AG2. This excess of ANT activity (in amplitude and/or in duration) and the second burst of the agonist (AG2) produce a small oscillation of the joint angle. Such small oscillations at the movement end were accepted as long as the error (MSE) calculated between the movement and its minimum-jerk profile was less than 1. Movements with too large end oscillation (see Figure 1F, right, profile (1)) were rejected.

In order to apply the definition of the triphasic command, we have used the electrical activity of muscles. However, since all models used in this study were based on non-spiking neurons (and hence non-spiking muscle), we used the non-spiking muscle activity to analyze triphasic command occurrences. In order to illustrate EMG profiles, for each model, we give examples of EMGs produced during a large movements (#80°), during a medium amplitude movement (#60°), and during a smaller movement (#45°) (see Figure 10C1-3).

**Figure 10:**
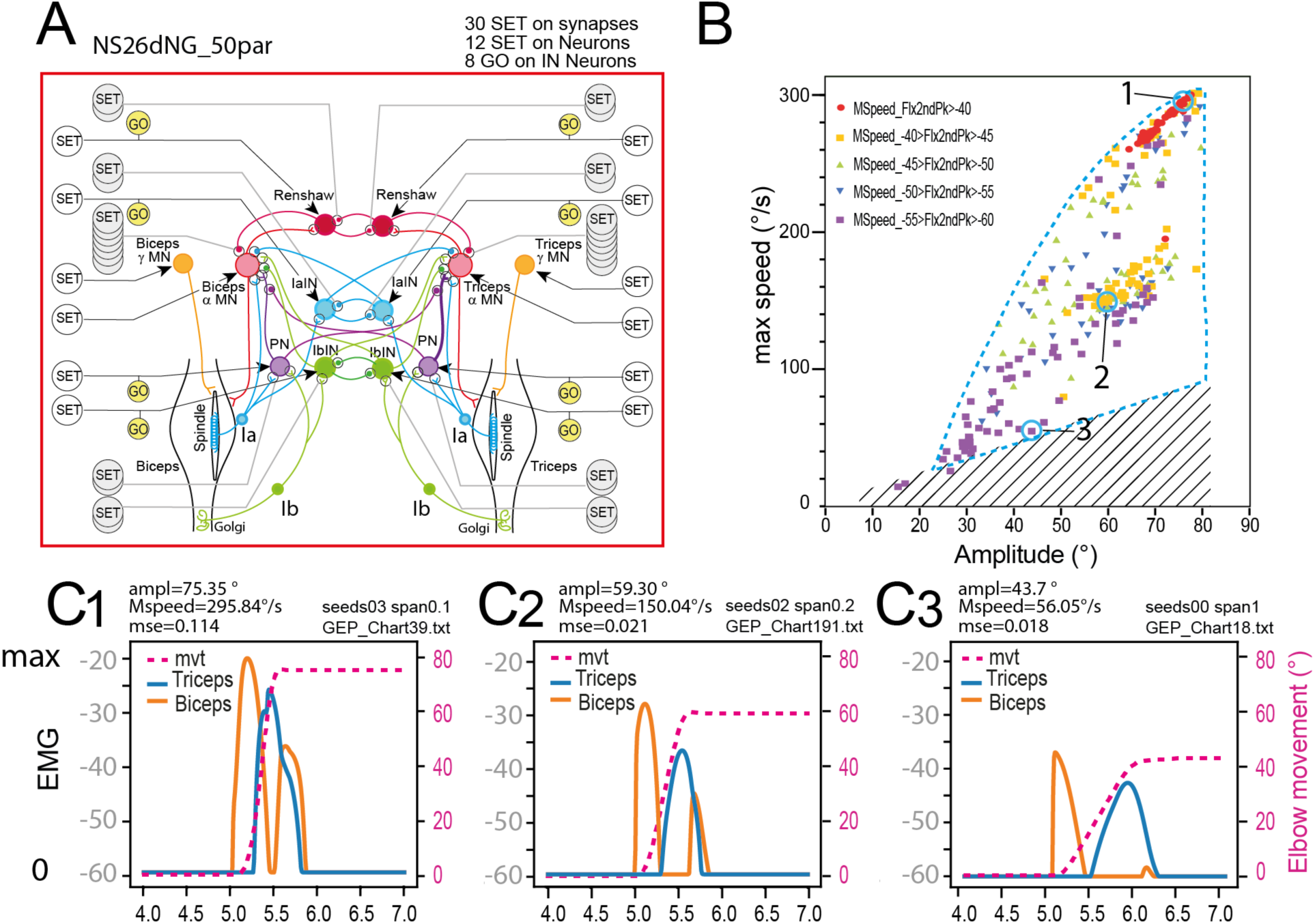
Presence of triphasic pattern in the behavior space with the complete 50 parameter model. **A**, Sensorimotor spinal network of the 50-parameters model. **B**, Presence triphasic pattern in the behavior domain uncovered by rGEP on this model. As can be seen triphasic pattern were found all over the domain (see blue dashed lines for the limits of the 50-parameter behavior domain). **C**, Examples of triphasic pattern for large (80°, C1), medium (60 °, C2) and small (45 °, C3) movements. These 3 examples are identified in B, with numbers 1,2 and 3.

#### 1) Triphasic commands with the complete (50-parameters) model

The occurrence of triphasic commands with AG1, ANT, AG2 peaks of activity were only observed with the **50-parameters model** (see the full network in Figure 10A). Triphasic patterns with two Flexor peaks, separated by a silent period during which an Ext peak was present, were observed in the whole domain of the valid behaviors produced by the 50 parameters model (Figure 10B). Three examples of triphasic pattern are given in Figure 10C1-3, for large movement (80°; Figure 10C1), medium movement (60°; Figure 10C2) and small movement (45°; Figure 10C3). It is interesting to note that for the largest movements the Ext EMG peak was narrow, and its width increased for smaller movements (compare orange trace in Figure 10C1-3). It is also interesting to note that all of these movements were very close to a minimum jerk profile (their MSE were 0.069, 0.036 and 0.012, respectively). Finally, in all of the valid movements with a triphasic pattern, the activity between the two AG peaks went back to 0.

Here we represented only the movements associated with a clear triphasic pattern. However, such movements associated with a true triphasic pattern represent only 5% of the valid movements found by rGEP. Indeed, in the great majority of movements found by rGEP, a single Flx EMG peak is followed by a single Ext EMG peak. This is generally enough to produce a perfect minimum jerk movement.

#### 2) Triphasic commands with the other models

In the other models, we could not find any valid movement with a triphasic pattern. Even in the 36-parameters model (complete model deprived of the 1b afferent and Ib Interneurons), there were valid movements very close to minimum jerk (MSE<0.05) but the second Flx EMG peak was always absent. Nevertheless, we could find very rare pseudo triphasic patterns. They were rejected either because the second peak was so small that it did not elicit any force in the Biceps, or because their profile did not meet the minimum jerk criterium (MSE >1). Moreover, all of these pseudo triphasic profiles were found in slow movements (data not shown).

#### 3) Elaboration of Triphasic commands in spinal sensorimotor networks

An example of triphasic commands with AG1, ANT, AG2 peaks of activity was used to examine the temporal relationships between EMG, Alpha MN activity, PN activity and sensory 1a and 1b activity (Figure 11). These neurons represent all the spinal sensorimotor neurons that participate to the dynamical activity leading to the triphasic command (Figure 11A, B). Only gamma MNs are not represented because they only display step activation. Constitutively, FlxEMG and ExtEMG are directly dependent on the activity of FlxAlpha and ExtAlpha, respectively (Figure 11C). In the same way, the activity of FlxAlpha and ExtAlpha should depend on FlxPN and ExtPN, respectively (see Figure 11A), but the time course of the peaks observed in Alpha MNs and their corresponding PN are different (Figure 11D). The difference is small for FlxPN, the peak of which is delayed compared to FlxAlpha peak, but larger for ExtPN, the peak of which is earlier than that of ExtAlpha. These mismatches can be explained because Alpha MNs receive also other input from their corresponding 1a afferents (Figure 11A). The role of Flx1a will be more precisely analyzed in the next paragraph. Note simply that Flx1a activity (yellow curve in Figure 11E) decreases during most of the FlxAlpha. By contrast the role of Ext1a is more directly understandable because Ext1a depolarizes and peaks at the same time as ExtAlpha (compare light-blue and green traces in Figure 11E). Interestingly, 1b afferent present peak activities that correspond to their corresponding Alpha MN activities (Figure 11F). For example, the large early peak of Flx1b contributes to the activation of FlxAlpha via its connection to FlxPN (compare pink and red traces in Figure 11F). Finally, the activity of Ext1b also contributes to the late part of ExtAlpha activation via its connection to ExtPN (see the second bump on ExtAlpha activity that corresponds to the peak of Ext1b and to the peak of ExtPN in Figure 11F).

**Figure 11:**
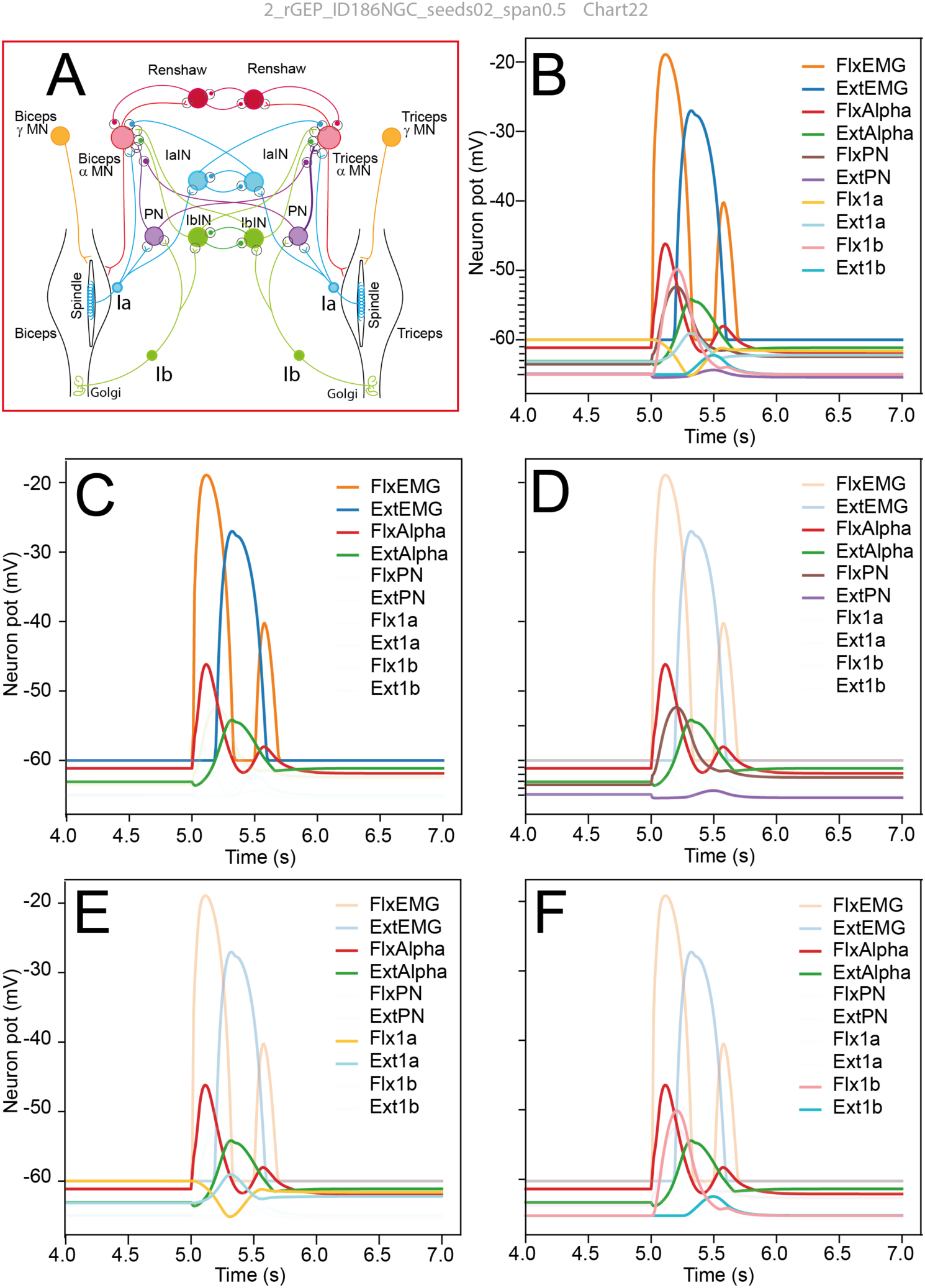
Activity of spinal neurons during triphasic pattern with the complete 50 parameter model. **A**, Sensorimotor spinal network of the 50-parameters model. **B**, Activity of Alpha MNs, PN interneurons, 1a and 1b sensory neurons during a triphasic command. **C**, Relationship between biceps EMG (FlxEMG), triceps EMG (ExtEMG), biceps alpha MN (FlxAlpha) and triceps alpha MN (ExtAlpha). **D**, Relationship between propriospinal interneurons (FlxPN and Ext PN) and their corresponding MNs (FlxAlpha and ExtAlpha). **E**, Relationship between 1a sensory afferents (Flx1a and Ext1a) and their corresponding MNs (FlxAlpha and ExtAlpha). **F**, Relationship between 1b sensory afferents (Flx1b and Ext1b) and their corresponding MNs (FlxAlpha and ExtAlpha).

To go further in the analysis of how triphasic pattern can be elaborated in spinal sensory-motor circuits, we have examined more precisely the impact of each sensory feedback, by replaying the same example of movement as presented in Figure 11, after suppression one of the four sensory feedback. The results are presented in Figure 12 for the movement launch, and Figure 13 for the movement stop.

**Figure 12:**
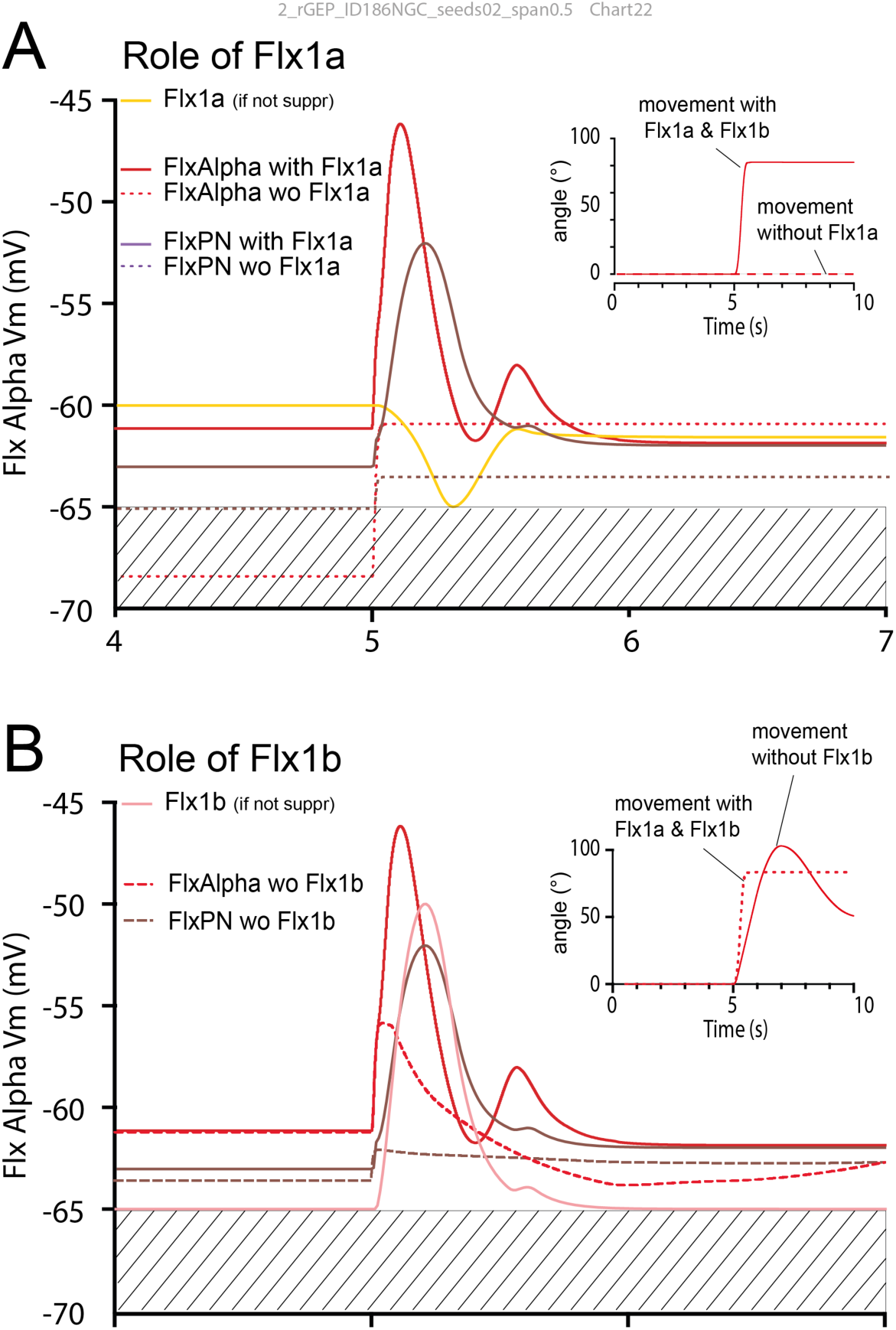
Role of Flx1a and Flx1b in onset of triphasic pattern. **A**, Role of Flx1a. The same movement as presented in Figure 11 was replayed with inactivation of the Flx1a afferent. The control FlxAlpha and FlxPN activities are presented in red and brown traces, respectively. The control Flx1a activity is presented in a yellow trace. After inactivation of the Flx1a afferent, no movement is produced (see dashed line in inset) **B**, Role of Flx1b. Same disposition as in **A**. The control Flx1b activity is presented in a pink trace After Flx1b afferent is inactivated, a slow movement is still produced (see dotted line in inset), but it is unable to stabilize. Flx1Alpha activity is also smaller (see red dashed line). Moreover, the FlxPN peak (full brown line) disappears when Flx1b is inactivated (dashed brow line).

**Figure 13:**
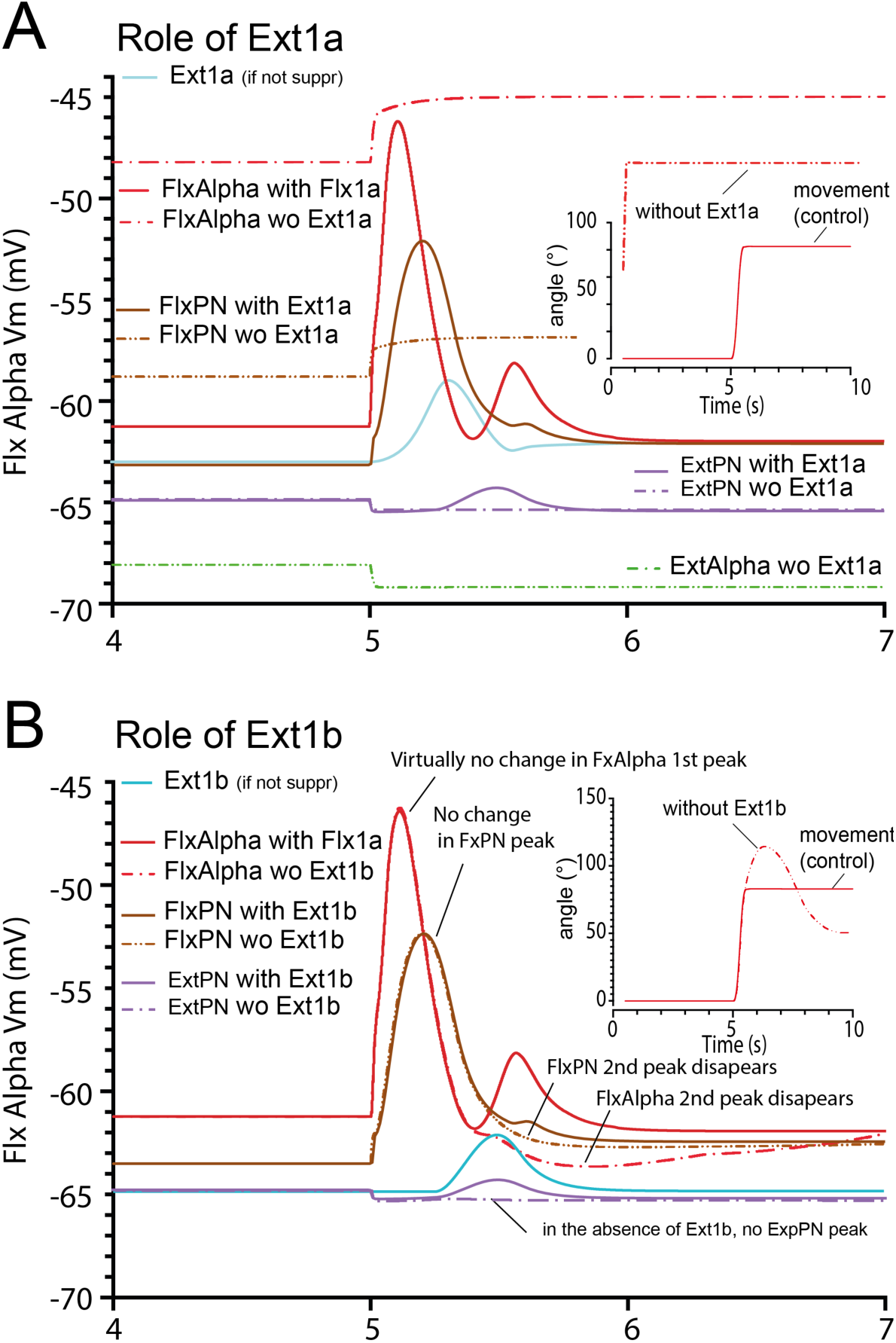
Role of Ext1a and Ext1b in shaping triphasic pattern. **A**, Effect of Ext1a inactivation. As in Figure 12, control FlxAlpha and FlxPN activities are presented in red and brown traces, respectively. The control Ext1a activity is presented in a lightBlue trace. After inactivation of the Ext1a afferent, movement starts at the onset of the preparatory phase (see dashed line in inset), and was therefore rejected. **B**, Role of Flx1b. Same disposition as in **A**. The control Ext1b activity is presented in a blue trace After Ext1b afferent was inactivated, a movement is still produced (see dotted line in inset), but with a huge overshoot. Moreover, the second FlxAlpha peak (full red curve), the second FlxPN peak (brown curve) and the ExtPN peak (violet curve) disappear when Flx1b is inactivated (dashed lines).

##### (i) Role of Flx1a in movement launch

When Flx1a was inactivated (i.e., its membrane potential maintained artificially constant at -65 mV), we observed that the membrane potential of FlxAlpha was hyperpolarized to -68 mV in the preparing phase (see dotted red trace in Figure 12A). Similarly, in the absence of Flx1a activity, FlxPN was also hyperpolarized to -65 mV during the preparatory phase (see dotted brown trace in Figure 12A). These results indicate that during the preparatory phase, in the normal configuration, Flx1a was continuously activating FlxAlpha (and FlxPN), but FlxAlpha did not fire because of the hyperpolarizing SET command it received. When the GO command is given at t=5s, FlxAlpha is immediately freed from this hyperpolarization and will be activated by Flx1a. Interestingly, this source of activation will be transient because as the forearm get flexed, the Flx1a signal decreases and rapidly vanishes (see red trace in Figure 12A).

##### (ii) Role of Flx1b in movement launch

When we replayed the parameters of the same movement presented in Figure 11 and Figure 12, after inactivating Flx1b, we still observed a movement, but it was slower (compare red trace with dotted red trace in Figure 12B inset). This result is in accordance with the reinforcing role of 1b afferent on their muscle during the onset of movement observed in physiology (Donelan & Pearson, 2004).

##### (iii) Role of Ext1a and Ext1b in movement stop

We then replayed the same movement parameters after inactivating Ext1a (Figure 13A). In the absence of Ext1a, the preparatory phase was too much disturbed (movement started at t=0s instead of t=5s) to conclude about the role of Ext1a in the movement stop. The role of Ext1a in the activation of ExtAlpha was however presented in Figure 11E and in the corresponding text (see above).

The role of Ext1b was easier to decipher, because in its absence, the second FlxAlpha peak totally vanishes (see dashed red curve in Figure 13B). The suppression of the 2^nd^ FlxAlpha peak is also linked with the disappearance of the 2^nd^ FlxPN peak (see dashed brown curve in Figure 13B). Moreover, the ExtPN peak also disappears (see violet curve in Figure 13B).

##### (iv) Which Ext1b pathway is involved in movement stop and 2^nd^ FlxAlpha peak in triphasic pattern?

From the result presented in Figure 13B, we conclude that Ext1b is involved in the FlxAlpha 2^nd^ peak of triphasic pattern. Then, we dissected out all pathways between Ext1b and FlxAlpha (Figure 14). If we consider all these pathways (Figure 14A), it is easy to replay the simulation of this movement, after selectively suppressing a single synapse to estimate its role in the transmission of Ext1b Feedback. When the Ext1bIN->FlxAlpha synapse was suppressed (blue dashed curve in Figure 14A2) the FlxAlpha second peak was only slightly decreased (see blue dashed curve in Figure 14B). When the ExtPN->FlxPNPre was suppressed (orange dashed line in Figure 14A3), the FlxAlpha second peak was unchanged (see orange dashed curve in Figure 14B). The same absence of effect was also observed when the Ext1bIN->Flx1bIN synapse was suppressed (orange dashed line in Figure 14A5 and in Figure 14B). A slight decrease in the second peak was also observed wen these three synapses were suppressed at the same time (red dashed curves in Figure 14A6 and Figure 13B).

**Figure 14:**
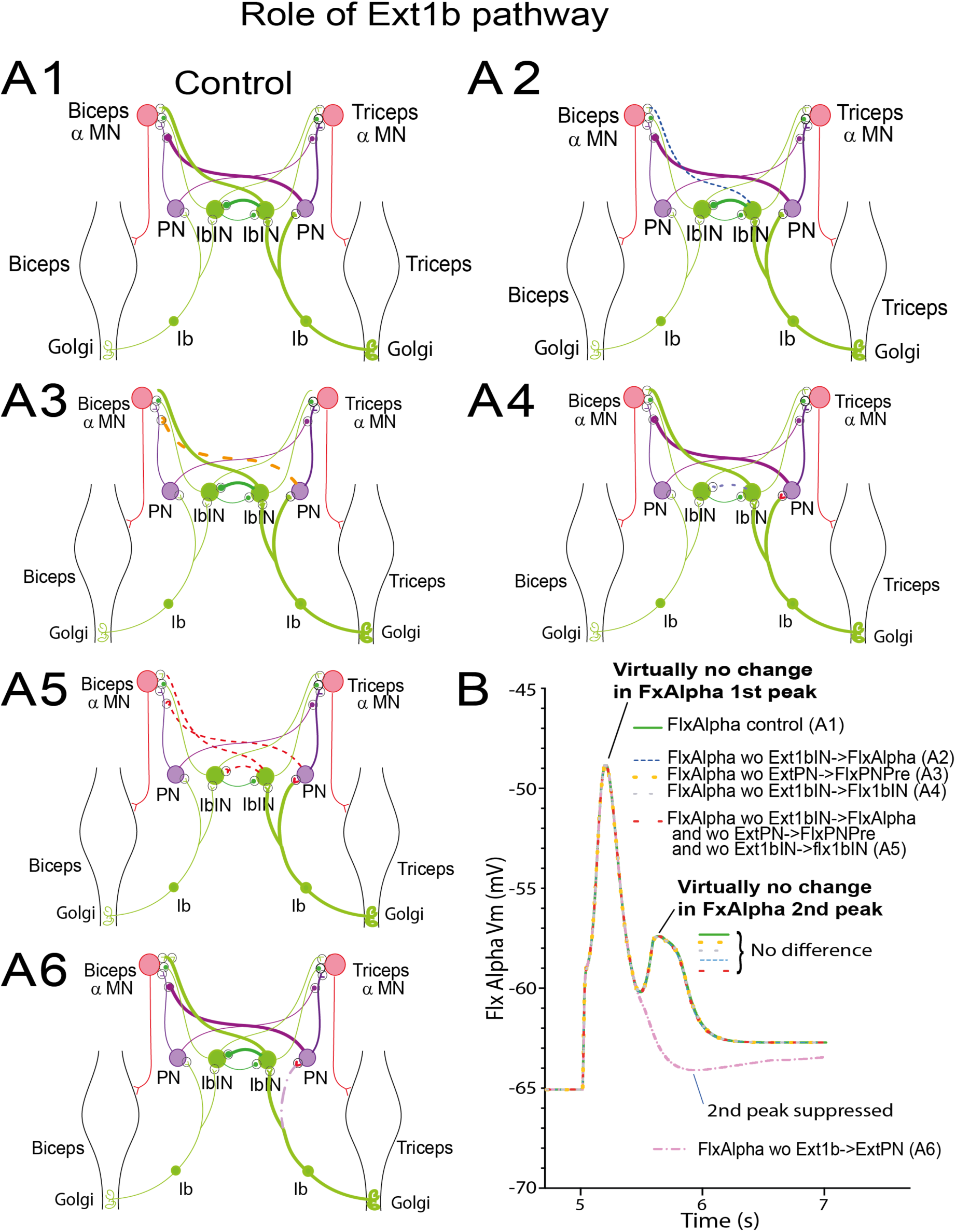
Role of Ext1b pathways in the production of the 2^nd^ FlxAlpha peak of triphasic pattern. **A**, Exploration of the various pathways from Ext1b to FlxAlpha (A1). Several replays of the simulation of the same movement as in Figure 12, 12 and 13) have been performed in which one of the involved synapses has been suppressed in each run (A2, A3, A4 and A5). In A6 the three synapses involved in crossing pathways have been suppressed at once. The result of the run of the model with thes diverse ablations are presented in B using the same color as the suppressed synapse. (see text for explanations).

By contrast, when the Ext1b->ExtPN synapse was suppressed (dashed violet curve in Figure 14A4), the second FlxAlpha peak totally disappeared (dashed violet curve in Figure 14B). These results indicate that the second FlxAlpha peak is not due to a cross connection from Ext1b to FlxAlpha. Therefore, if this 2ndFlxAlpha peak requires Ext1b feedback to be present, its effect is not central but rather is a consequence of strengthening of ExtAlpha by the Ext1b->ExtPN->ExtAlpha pathways.

This hypothesis was then tested in Figure 15 in which we assessed the consequence of suppressing the Ext1b->ExtPN synapse, onto FlxAlpha (red curves in Figure 15A), ExtAlpha (green curves in Figure 15A) and Ext1b activity (blue curves in Figure 15A). As can been observed, the suppression of the Ext1b->ExtPN synapse not only suppresses the FlxAlpha 2^nd^ peak, but also alters ExtAlpha activity profile and reduces the Ext1b peak itself. These later effects are produced via the mechanical system, because Ext1b->ExtPN synapse is excitatory as is the ExtPN->ExtAlpha synapse. Therefore, Ext1b activity contributes to reinforcing the ExtAlpha peak. Note that because of the positive feedback of Ext1b onto ExtAlpha, the increase of strength will in turn increase the Ext1b activity (compare Ext1b activity with and without Ext1b->ExtPN synapse in blue and dashed blue curves in Figure 15A). This effect strengthens the braking action he ExtAlpha on the flexion movement that will end more rapidly. This rapid change in movement speed is then perceived by Flx1a as can be observed in Figure 15B (see yellow curves). In the absence of Ext1b->ExtPN synapse, due to the lack of brake, speed decreases less rapidly and the Flx1a activity is reduced in its last part (after t=5.3s, see yellow dashed curve in Figure 15B). More importantly, the second Flx1a peak is also suppressed.

**Figure 15:**
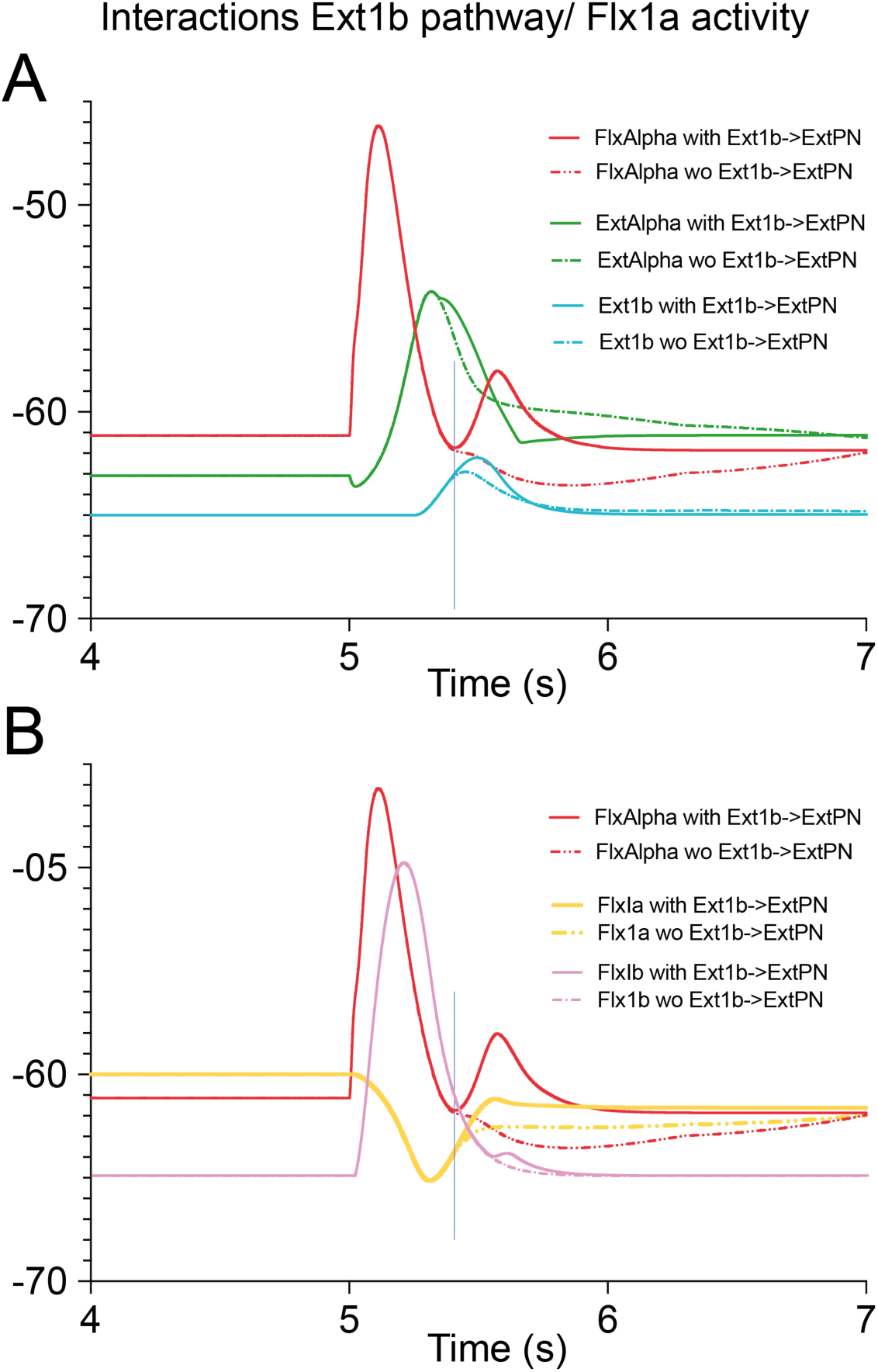
Interaction of Ext1b pathways and Flx1a activity. **A**, Contribution of Ext1b->ExtPN synapse on ExtAlpha activity and Ext1b feedback. **B**, Contribution of Ext1b->ExtPN synapse on Flx1a and Flx1b feedback. The FlxAlpha activity is presented in both graphs (A and B) to indicate the timing of the second peak.

The rapid change in movement speed is also perceived by Flx1b as can be observed in Figure 15B (see pink curves): this effect can be observed as a second peak on the Flx1b activity (plain pink curve in Figure 15B), that disappears when Ext1b->ExtPN synapse is suppressed (dashed pink curve in Figure 15B). The presence of these two sensory 2^nd^ peaks of Flx1a and Flx1b are responsible for the FlxAlpha 2^nd^ peak via direct connection and via FlxPN, respectively (see FlxPN 2ndPeak brown trace in Figure 13B).

##### (v) Confirmation of triphasic elaboration process in spinal circuits: role of 1a and 1b feedbacks?

In order to confirm the determinant role of Ext1b feedback in enhancing both ExtPN peak and brake action on flexion movement without role of crossing pathways, we have built a simplified version containing 1a and 1b feedbacks on PN and MNs, with no crossing pathways, no 1aINs and 1bINs, and no Renshaw INs. This simplified 16 parameters model is presented in Figure 16A.

**Figure 16:**
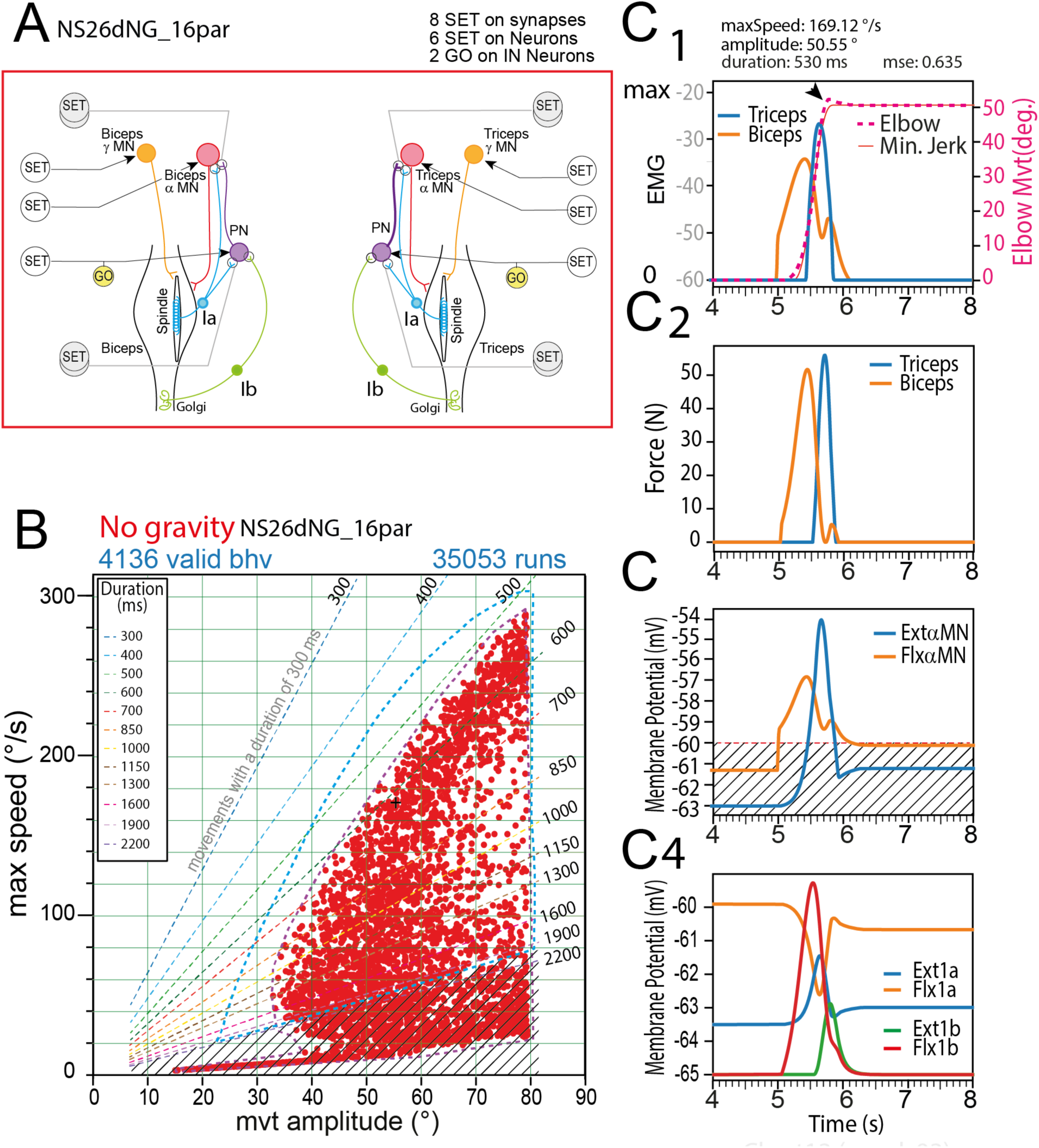
Minimal model with 1a and 1b feedbacks and no cross connections. **A**, Presentation of the 16-parameters model. **B**, Behavior space obtained with the 16-parameters model. Note that this behavior domain is almost as large as the one of the complete 50-parameters model (see blue dashed line). **C**, representative example of a valid movement (scale presented on the right side); EMGs from Biceps and Triceps (orange and blue traces, respectively) display a triphasic command; **C1**, Time course of the flexion movement with an amplitude of 50.5° and a maximal speed of 169.12 °/s; **C2**, Time course of muscle forces for the biceps (orange trace) and the triceps (blue trace). Note the two peaks of Biceps force; **C3**, Flexor and extensor MNs activities during the movement (the stripped domain represents subthreshold activity); **C4**, Muscle spindle and Golgi Tendon Organ feedbacks.

As can be seen, the behavior domain of this simplified 16-parameters model is almost as large as that of the complete 50-parameters model (Figure 16B). A representative valid behavior displaying a triphasic command is presented in Figure 16C. Note that on this example, movement presents a small overshoot (arrowhead in Figure 16C1). Note also that in this rapid movement the two flexor EMG peaks partially overlap, as was observed in physiological reports for rapid movements (see Figure1C-D in Brown and Cooke, 1981). Valid behaviors presenting a typical triphasic command, were found in most of the behavior domain of the 16-parameters model (Figure 17B).

**Figure 17:**
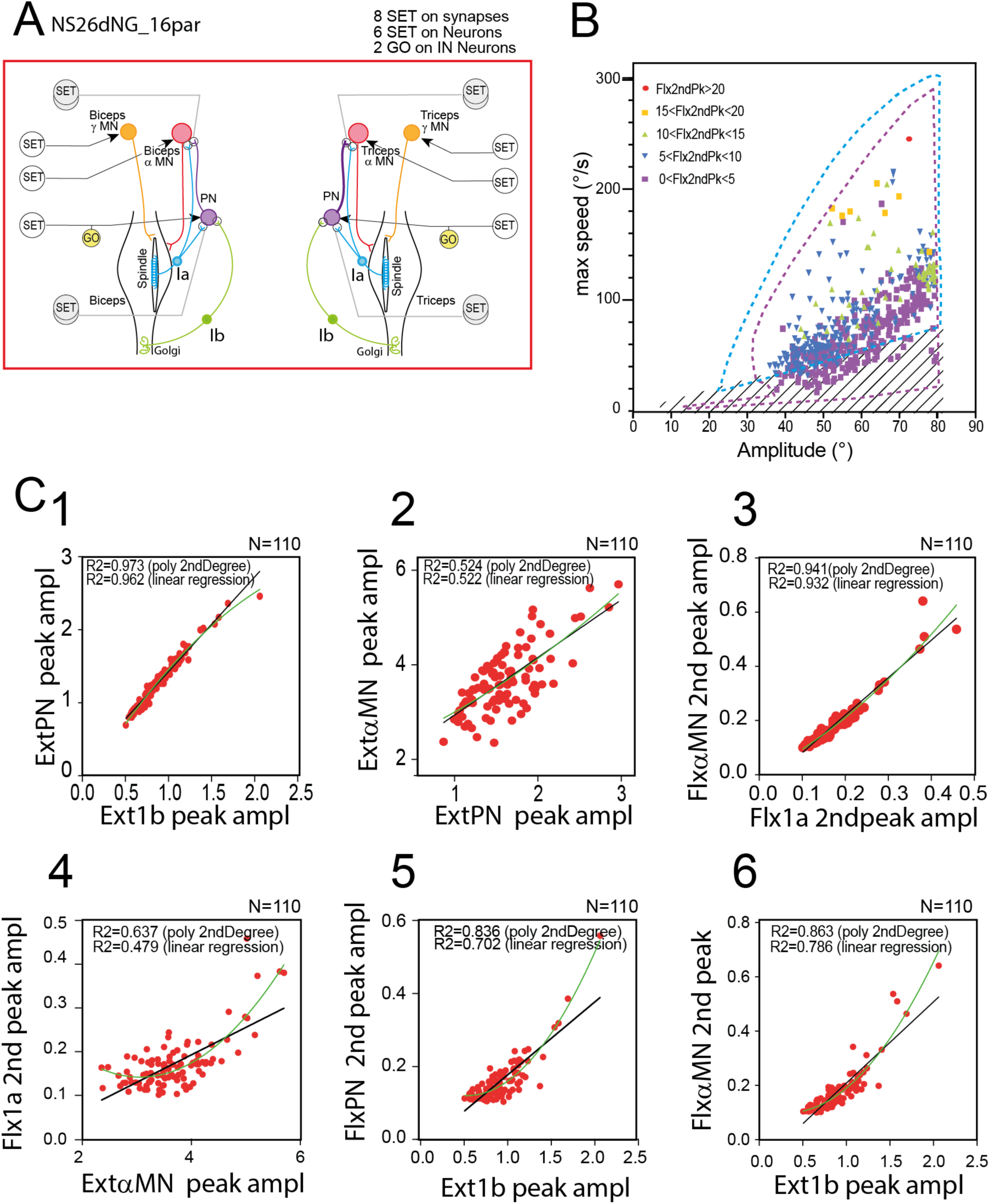
Analysis of triphasic patterns in the 16-parameters model. **A**, 16-parameters model circuits **B**, Repartition of triphasic pattern in the behavior space obtained with the 16-parameters model. Each occurrence is presented with a symbol indicating the amplitude of the 2^nd^ FlxMN peak: ranges are [0, 5] (violet squares), [5, 10] (blue triangles), [10, 15] (green triangles), [15, 20] (yellow squares) and >20 mV (red circle). **C**, Correlations between the amplitudes of Ext1b peak and ExtPN Peak (**C1**), ExtPN peak and ExtAlpha MN peak (**C2**), Flx1a 2^nd^ peak and FlxAlpha 2^nd^ peak (**C3**). These correlations indicate the strength of the functional synaptic links between the involved neurons. Correlations also exist between neuron activities involving a mechanical link (via proprioceptive feedback (**C4-C5**). For example, the amplitude of the ExtAlpha MN peak (indicating the brake power) is correlated to the amplitude of the Flx1a 2^nd^ peak (**C4**). Similarly, the amplitude of the Ex1b peak is correlated both to the amplitude of the FlxPN 2^nd^ peak (**C5**) and FlxAlpha 2^nd^ Peak (**C6**).

The behavior domain presented in Figure 16B was issued from a single seed. This choice was made to obtain the largest behavior domain based on the same strategy (see discussion). The presence of a single strategy in the whole behavior domain greatly facilitated the unravelling of the links between the various actors presented in Figures 13, 14, 15. These links were established by correlation analysis made on peak amplitudes of the various actors involved in the elaboration of triphasic commands (Figure 17C). All the links were thereby validated by significant correlation coefficients. As expected, Ext1b peak is linked to ExtPN peak (Figure 17C1, R^2^=0.962, p<0.00001), which is itself linked to the ExtAlpha peak (Figure 17C2, R^2^=0.522, p<0.00001). Similarly, the FlxAlpha 2^nd^ peak is linked to the Flx1a 2^nd^ Peak (Figure 17C3, R^2^=0.932, p<0.00001). All these links are supported by synaptic connections, but other correlations are the consequence of a mechanical link, via proprioceptive feedback (Figure 17 C4, C5 and C6): the amplitude of ExtAlpha peak is correlated to the amplitude of Flx1a 2^nd^ peak (Figure 17C4, R^2^ = 0.479, p<0.00001); the amplitude of Ext1b peak is correlated both to the amplitudes of FlxPN 2^nd^ peak (Figure 17C5, R^2^ = 0.702, p<0.00001) and FlxAlpha 2^nd^ peak (Figure 17C6, R^2^ = 0.786, p<0.00001). In addition, from this last graph, it is clear that when Ext1b peak amplitude reaches 0, FlxApha 2^nd^ peak also vanishes, thereby confirming the result of Ext1b “numeric ablation” (Figure 13B, 13B and Figure 15).

## Discussion

In the present modelling work, we explored the capacities of spinal sensorimotor networks to produce triphasic EMG pattern when activated by simple SET and GO step commands. We observed spontaneously occurring triphasic pattern EMGs produced by spinal sensorimotor circuits (50-parameters model) during the goal exploration process used to establish the behavioral capabilities of the system. Critically, the timing of such triphasic patterns were not predetermined in the simple step functions controlling their state. Particularly, the GO command, initiated the movement (t=5s) but did not give any indication for the observed triphasic EMG patterns. It is important to note that such triphasic pattern were observed in the whole range of the behavior domain (Figure 10B). **Therefore, triphasic pattern production for any movement seems to be an intrinsic property of sensorimotor spinal network in interaction with musculoskeletal system via MN activities and proprioceptive feedback**. Since the 50-parameters model involved both Ia spindle and GTO afferent feedback, we have used a dissecting out approach to evaluate the role of Ia and Ib afferents and their respective sensorimotor circuits in the elaboration of the triphasic commands.

### A. Necessity of spindle and GTO feedbacks in triphasic pattern

Removing Ia sensory feedback totally prevented the occurrence of any valid behavior (Figure 8B), indicating that the inherent capacity of spinal sensorimotor circuits to elaborate valid behaviors when interacting with a musculoskeletal system, requires spindle afferent signals.

By contrast, removing Ib sensory feedback and Ib sensorimotor circuits did not prevent spinal sensorimotor circuits to produce minimum jerk movements (Figure 6B). However, in the absence of GTO feedback, the extend of the valid behavior’s domain was moderately reduced (compare Figure 6B and Figure 5B). More strikingly, the absence of GTO feedback totally prevented the occurrence of triphasic commands, and indicates that although Ia feedback and Ia sensorimotor spinal circuits can handle the elaboration of minimum jerk movements, Ib afferent signals and Ib sensorimotor circuits are necessary (in addition to 1a sensory-motor circuits) to produce triphasic commands in our model. Note, however, that spinal reciprocal connections in sensorimotor circuits, seem to be important for fast valid movements to be produced (compare Figure 5B with Figure 16B). It seems that although valid movements with triphasic command in the same range of amplitudes can be produced in the absence of reciprocal connections (Figure 16B), for each amplitude, the maximal speed is significantly reduced. Yet, it is important to stress that reciprocal connections via spinal interneurons (Renshaw, Ia, Ib, PN) are not sufficient on their own to produce valid movements, i.e., in the absence of proprioceptive feedback (Figure 9B). Therefore, this study demonstrates that spinal sensorimotor networks fed by spindle afferent signals and GTO afferent signals, have the capacity to handle the production of triphasic command pattern in the whole range of movements of the behavior space.

It is then highly likely that spindle and GTO afferents participate in the elaboration of this triphasic command in animals and human. But if this conclusion is true, how to explain the finding that triphasic patterns were still produced in transient deafferentation experiments (Sanes & Jennings, 1984)? In fact, this experiment does not totally invalidate the **participation of sensorimotor circuits** in the shaping of the triphasic command, because the ischemic block affected the afferent activity of the muscles below the elbow, leaving intact both the efferent copy of their motoneurons via Renshaw’s interneurons and all proprioceptive feedbacks from all proximal muscles. The widespread extent of these feedback circuits (Cavallari *et al*., 1992), and the strong modulation of proximal muscles to stabilize posture during distal phasic movements, were not fully appreciated until later (Massion, 1992).

### B. Role of spindle feedback in triphasic pattern

We have shown that 1a feedback is essential for the spinal sensorimotor circuits to handle valid movements. Indeed, it is interesting to address the question of how simple step commands can elicit dynamic responses in motor activities, i.e. transient activation of flexor at two times ((i) for launching the movement, (ii) just before stabilization at the end of the movement), and transient activation in the extensor to stop the movement.

Let’s start with the first Flex transient activation. When we look at the result of suppressing Flx1a feedback (Figure 12A), we had noted that without Flx1a feedback the movement did not start, because the strategy uncovered by GEP consisted in a strong enhancement of Flx1a sensitivity to maximize its discharge during the preparatory phase. To avoid this sensory activity to activate FlxAlpha MN, the SET command sent a hyperpolarizing order to FlxAlpha MN. When the GO command was sent to FlxAlpha MN, this hyperpolarising action was suppressed and the Flx1a sensory feedback could strongly activate FlxAlpha MN. But, because movement was launched, the biceps muscle started to shorten and Flx1a signal decreased (see yellow curve in Figure 12A). This is how the step command elicited a transient activation of FlxAlpha MN, via sensory coding of muscle length (i.e. via the movement itself).

The transient activation of ExtAlpha MN (Figure 11E, green curve) is also directly related to Ext1a feedback (Figure 11E, blue curve). But its transient nature is related to the **dynamic response of Ext1a stretch** (i.e., to the speed of the stretch – see Figure 4A). Note that a possible improvement of our model of spinal sensorimotor circuit would be to introduce both dynamic and static gamma commands. This would improve handling of 1a feedback transients. This was not possible in AnimatLab but future work should address this question.

### C. Role of GTO feedback in triphasic pattern

It seems that 1b feedback is necessary for occurrence of triphasic command, but not sufficient because it cannot produce triphasic command without spindle feedback. So, how 1b feedback can help 1a afferent feedback for triphasic pattern production in spinal sensorimotor circuits? If we look at the activity of FlxPN (Figure 11D) and Flx1b (Figure 11F) in the 50 parameters model, we note that they display similar activity along time course. This indicates that Flx1b signals play a great role in assisting the Flx activity at movement onset. When Ib feedback is suppressed, maximal speed is reduced (see inset in Figure 12B). More importantly, the transient FlxAlpha peak totally disappears (compare red dashed line and red curve in Figure 12B). This means that Flx1b and Flx1a activities are both necessary to launch the movement. Then while the muscle shortens, Flx1a activity decreases rapidly, and because the force produced by the Biceps muscle decreases due to (i) muscle shortening and (ii) to FlxAlpha MN actvity decrease (consecutive to Flx1a decrease), the activity of Flx1b becomes transient. This involvement in force enhancement was also described in physiological conditions in cat (Donelan & Pearson, 2004). It seems that 1b contribution consists mainly in this enhancement in the flexor phase of the movement (i.e., movement initiation), but 1b feedback seems also involved in reinforcing ExtAlpha MN activity when the extensor is stretched during the flexion movement (compare ExtAlpha activity with and without Ext1b feedback in Figure 13B) and certainly contributes to shortening the brake phase. Therefore, in the absence of 1b feedback, the brake is less powerful and this phase is too slow for a minimum jerk profile (compare end of movements in the complete model – Figure 5C1, with the model derived of 1b feedback – black arrow in Figure 6C1). Ext1bpeak induces a rapid slowdown at the end of the movement, which is responsible for the second peak of Flx1a activity (due to **velocity and acceleration sensitivity of 1a afferents** - see yellow trace in Figure 11E and Figure 12A). Consequently, this Flx1a 2^nd^ peak, in turn activates the Flx alpha MN (red trace in Figure 12A). In addition, Flx1b also sense this rapid brake and produces a second peak (see pink trace in Figure 12B) that is transmitted to FlxPN (see brown trace in Figure 12 A, B). Finally both Flx1a 2^nd^ peak and FlxPN 2^nd^ peak converge onto FlxAlpha MN (see yellow, brown and red trace in Figure 12A).

### D. Role of descending commands and spinal networks in triphasic pattern

The fact that Ia spindle and Ib feedbacks and their spinal sensorimotor circuits are capable of handling triphasic commands all over the behavior domain, does not exclude, however, that in physiological conditions, descending command also participate in the elaboration of triphasic commands. Recordings from M1 motor cortex indicate that unit and global activities can be correlated either with static and dynamic forces and torques around single joints (Cabel *et al*., 2001), or with muscle synergies (Holdefer & Miller, 2002). However, the question of the origin of triphasic activities in M1 is still open because correlation is not causation, and triphasic activities in M1 could simply result from efference copy from spinal interneurons and afferent feedback from proprioceptors. Whatever the origin of M1 triphasic activity, it is likely that such precise descending command interact with spinal sensorimotor networks. Future research in this domain will have to unravel how complex descending commands interact with spinal sensorimotor process that can handle by itself triphasic commands.

In the present report, we used simple step commands to control spinal sensorimotor circuits. Similarly, some authors have argued that because EMGs are organized in triphasic pattern with silent periods, this indicated that descending commands did not operate in a smooth and continuous manner **(Leib *et al*., 2020)**. Indeed, the simple model (bang-bang optimal control model) proposed by these authors is more structured in time than the simple step functions we used here. In the bang-bang control, the control signal abruptly switches between a maximum and a minimum value at certain sparse points in time. The control function is made of three constant segments (with two transitions). The bang-bang model explains both abrupt changes in muscles activity (triphasic command) and the smooth reaching trajectory. Interestingly, rGEP was capable of uncovering a set of valid behaviors (minimum jerk movements) relying incidentally on triphasic commands (with abrupt changes in activity), although this triphasic command was not explicitly searched for nor included in the process with simple step commands. Therefore, it is possible that spinal network properties (and the associated biomechanical effector system) possess the capacities of handling such complex dynamic commands just by giving the correct level of stimulation of target spinal neurons and precise synaptic gain settings. In a more general perspective, these spinal sensorimotor circuits could be considered as the physiological support of movement primitives proposed by Degallier and Ijspeert (2010) that can be handled by simple step commands from the brain. In this case, the motor primitives not only encompass movement’ start dynamics but also movement’ stop dynamics with a unique step command.

### E. Triphasic pattern and minimum jerk

Simulation studies have suggested that minimum-jerk models (Flash & Hogan, 1985) and minimum torque-change models (Uno *et al*., 1989) were likely not capable to predict muscle actions (Osu *et al*., 2004) and thus triphasic pattern. Here, we used a different approach that involved a musculoskeletal model associated to spinal sensorimotor networks, and a search method (rGEP) to revisit the capability of spinal sensorimotor circuits to produce triphasic pattern of muscle activation.

Indeed, rGEP uncovered a great diversity of parameters associated with a given valid movement. Among these parameters, some will result in triphasic commands. This solution was found all over the behavior domain of the model. But other parameter sets do not fit the definition of triphasic command. For example, although the silence of the agonist between the two agonist peaks (AG1 and AG2) was found in most of the 50-parameters model valid movements (Figure 10B), there were also solutions with no 2^nd^ Flexor peak all over the behavior domain of this complete model.

The fact that we imposed a minimum jerk profile for movements had consequences on the type of triphasic command retained by rGEP. Indeed, when we consider fast movements that were retained to define the triphasic pattern (Hallett *et al*., 1975), a small oscillation at the end of the movement is often present. The origin of this final oscillation comes from ANT peak that was slightly too strong, resulting in a final oscillation that required the second agonist peak (AG2) to be damped (Berardelli *et al*., 1996). The minimum jerk profile virtually imposes the absence of such oscillation at the end of the movement, and thereby reduces dramatically the amplitude of AG2.

### F. Limits and perspectives of the study

#### No separate Static and Dynamic gamma MNs

All models of the present study used a mixed type of gamma MN (one for the biceps and one for the triceps). Ideally, we should have used two types of gamma MNs for each muscle: a static gamma MN and a dynamic gamma MN, as was the case in the model of spinal-like regulator (Raphael *et al*., 2010), but these distinct gamma control does not exist at present in AnimatLab. However, the activation of the gamma MN of our model increased spindle sensitivity to both position and velocity, and this single type of gamma MN was enough to produce the desired behavior of the model under rGEP. Certainly, the possibility for the step commands to act differently on spindle length coding and speed/acceleration coding feedbacks should benefit to the possibilities of control (see “Role of spindle feedback in triphasic pattern” above), and likely extend the behavior domain of the model. This point will need future work to estimate the changes brought by the two gamma MN types.

#### Absence of gravity

We presented here the results of movements conducted in the absence of gravity. The presence of gravity during elbow flexion exerts an increasing torque that opposes the action of the biceps and could potentially obscure the role of the triceps in slowing down the movement, particularly when the movements are not sufficiently fast. Note that in many experiments the effects of gravity are eliminated by performing movements on a frictionless surface, in the horizontal plane. The present results also apply to this experimental condition.

It will be interesting in future investigations to add gravity to see how triphasic pattern evolve, depending on movement trajectory, and how spinal network can handle it. Such studies should show smaller ANT burst when gravity is present and tend to slow down the movement as the forearm becomes more horizontal during flexion movements. Inversely, if movements start from a flexed position (say 110° in our model), and stops at 20°, ANT burst should be dramatically increased and thus AG2 burst should increase accordingly.

#### Simple flexion movement

In the present study, we used a single joint (elbow) and movement consisted in elbow flexion. This was made because minimum jerk is more adapted to this situation (Nakano *et al*., 1999). It would be interesting to generalize the results of the present study by using more complex models including two joints, and mono- and bi-articular muscles, and considering endpoint reaching movements as was used in Tsianos et al. (2014).

#### GEP and exploration of redundancy

The systematic investigation of the behavioral constraints of reduced spinal circuitry highlights the functionality of a system that might otherwise be considered excessively redundant. For organisms navigating real-world environments, it may be more important to get the largest possible range of performances from their musculoskeletal systems rather than concentrating on the narrow range of tasks typically examined in laboratory settings. In this line of thought, a potential development of the present work would be to evaluate the adaptative role of redundancy by introducing unexpected perturbations or constraints, and see how redundancy helps the system to find appropriate solutions. Notably, such research would highlight the importance of spinal networks in facilitating rapid adaptation under the brain’s supervision as is the case in reservoir computing (Tanaka *et al*., 2019).

Finally, the present modelling study offers an opportunity to test the physiological roles of Ia and Ib proprioceptive feedback in voluntary movements. Since the model pointed out the specific roles of Ia and Ib at specific timings, it should be possible to test if the involved circuits are responsible for the elaboration of the triphasic commands observed during rapid arm flexion, for example by delivering perturbations at these specific time windows and see if muscle responses are in accordance with similar perturbations delivered in the model.

## Author Contributions

**Conceptualization:** Daniel Cattaert, Aymar de Rugy, Pierre-Yves Oudeyer.

## Funding acquisition

**Methodology:** Daniel Cattaert, Bryce Chung, Florent Paclet, Pierre-Yves Oudeyer, Aymar de Rugy.

**Project administration:** Aymar de Rugy.

**Investigation:** Daniel Cattaert, Matthieu Guemann.

**Software:** Daniel Cattaert, Bryce Chung, Florent Paclet.

**Supervision:** Aymar de Rugy.

**Validation:** Daniel Cattaert, Aymar de Rugy, Pierre-Yves Oudeyer.

**Writing – original draft:** Daniel Cattaert.

**Writing – review & editing:** Daniel Cattaert, Matthieu Guemann, Florent Paclet, Bryce Chung, Pierre-Yves Oudeyer, Aymar de Rugy

## Acknowledgments

This work was supported by The Centre National de la Recherche Scientifique (CNRS) and the University of Bordeaux.

